# Structural basis for the context-specific action of classic peptidyl transferase inhibitors

**DOI:** 10.1101/2021.06.17.448903

**Authors:** Egor A. Syroegin, Laurin Flemmich, Dorota Klepacki, Nora Vazquez-Laslop, Ronald Micura, Yury S. Polikanov

## Abstract

Ribosome-targeting antibiotics serve both as powerful antimicrobials and as tools for studying the ribosome. The ribosomal catalytic site, the peptidyl transferase center (PTC), is targeted by a large number of various drugs. The classical and best-studied PTC-acting antibiotic chloramphenicol, as well as the newest clinically significant linezolid, were considered indiscriminate inhibitors of every round of peptide bond formation, presumably inhibiting protein synthesis by stalling ribosomes at every codon of every gene being translated. However, it was recently discovered that chloramphenicol or linezolid, and many other PTC-targeting drugs, preferentially arrest translation when the ribosome needs to polymerize particular amino acid sequences. The molecular mechanisms and structural bases that underlie this phenomenon of context-specific action of even the most basic ribosomal antibiotics, such as chloramphenicol, are unknown. Here we present high-resolution structures of ribosomal complexes, with or without chloramphenicol, carrying specific nascent peptides that support or negate the drug action. Our data suggest that specific amino acids in the nascent chains directly modulate the antibiotic affinity to the ribosome by either establishing specific interactions with the drug molecule or obstructing its placement in the binding site. The model that emerged from our studies rationalizes the critical importance of the penultimate residue of a growing peptide for the ability of the drug to stall translation and provides the first atomic-level understanding of context specificity of antibiotics that inhibit protein synthesis by acting upon the PTC.

## INTRODUCTION

Protein synthesis is catalyzed by the ribosome, one of the most conserved and sophisticated molecular machines of the cell. The ribosome is a preferred antibiotic target and many classes of drugs selectively bind to the functional centers of the bacterial ribosome and prevent cell growth by interfering with different aspects of protein synthesis. Ribosome-targeting antibiotics are indispensable both as therapeutic agents and as tools for basic research. For example, the identity of the key ribosome functional centers and the modes of their operation have been established using antibiotics as structural and functional probes^1^. Understanding the true molecular mechanisms of action of the newest as well as the “classic” ribosome inhibitors is critical not only for our ability to develop new potent antibacterials but also to expand our knowledge about the fundamental principles of ribosome functioning. In most cases, ribosomal antibiotics interfere with protein synthesis by binding at various functional centers of the ribosome and either lock a particular conformation of the ribosome or hinder the binding of its ligands. The catalytic peptidyl transferase center (PTC) located at the heart of the large ribosomal subunit is the site targeted by the broadest array of inhibitors belonging to several distinct chemical classes, including phenicols, lincosamides, oxazolidinones, pleuromutilins, streptogramins, macrolides, and others.

One of the oldest (and hence “classic”) and the best-studied PTC-targeting antibiotics is chloramphenicol (CHL). Although CHL specifically targets bacterial 70S ribosomes, it can also bind to mammalian mitochondrial ribosomes^2-4^, causing major side effects^5,6^. As a result, the clinical usage of CHL is currently limited to developing countries, where it is used as an affordable alternative to the more expensive antibiotics. Nevertheless, other drugs of this class, such as florfenicol^7^, are widely used in the United States to treat infections in farm animals. However, despite CHL being studied for decades, we still lack a full understanding of the very basic principles underlying either the toxicity or the mode of the drug action for this class.

Multiple structures of CHL bound to vacant ribosomes from different bacterial species located its binding site in the A site of the PTC^8-10^. This observation resulted in the “classic enzymology”-inspired idea that the drug molecule acts simply as a competitive inhibitor^11^ that prevents either binding or proper placement of the aminoacyl moiety of any incoming aminoacyl-tRNA (aa-tRNA) into the ribosomal A site, resulting in inhibition of peptide bond formation. However, this commonly accepted model of CHL action fails to explain several experimental observations, such as the differential inhibition of translation of specific mRNA templates^12,13^ or only partial inhibition of puromycin-mediated release of nascent chains in polysomes even at saturating CHL concentrations^14^. The perceived ability of CHL to indiscriminately inhibit the formation of any peptide bond also conflicts with its role as an inducer of resistance genes. Activation of a number of CHL resistance genes relies on the antibiotic-promoted arrest of translation at specific codons within upstream regulatory leader open reading frame (ORF)^15,16^. Thus, to activate the expression of the resistance locus in response to the antibiotic assault, the ribosome should be able to progress through several leader ORF codons to reach the site of the programmed translation arrest. Therefore, it had remained unclear how the ribosome could polymerize a segment of the leader peptide if CHL indiscriminately inhibits the formation of any peptide bond. Detailed analysis of ribosome profiling data of the action of CHL in the bacterial cell^17^, as well as *in vitro* primer extension inhibition (toe-printing) assays^17^ and single-molecule Förster resonance energy transfer (smFRET)^18^ approaches on several representative ORFs, provided clear evidence that CHL does not block the formation of every peptide bond with the same efficiency but instead interferes with translation in a context-specific manner. The nature of specific C-terminal residues of the nascent peptide, as well as of the A-site acceptor, was found to strongly influence the ability of CHL to inhibit peptidyl transfer. The presence of alanine, and to a lesser extent of serine and threonine, in the penultimate position of the peptide is conducive to CHL-induced ribosome stalling^17,18^. In contrast, glycine in the P or A sites of the PTC strongly counteracts the inhibitory effect of CHL^17,18^. Unfortunately, all of the available structures of ribosomes associated with CHL show how the antibiotic molecule binds to the PTC of only either vacant ribosomes^8,9^ or ribosomes in complex with deacylated tRNA substrates^10^, and therefore could not provide structural insights into the observed context-specific activity. The key missing piece of information that prevents understanding the molecular mechanisms of the context specificity of PTC-targeting antibiotics is the lack of structures showing how the action of the drug, bound in the A site of the ribosomal catalytic center is affected by the nature of both, the donor (peptidyl-tRNA) and acceptor (aa-tRNA) substrates. Such structures would closely mimic the state of the ribosome in a living cell at the moment of the encounter with the drug and would provide the most accurate information of recognition of the translating ribosome by the antibiotic.

In this work, we solved for the first time high-resolution structures of the ribosome in the functional state during its interaction with the PTC-targeting antibiotic CHL. We present the structures of *Thermus thermophilus* (*Tth*) 70S ribosomes containing combinations of peptidyl-tRNA and aa-tRNA analogs that are either conducive to the drug action or render the peptide bond formation reaction immune to the presence of the antibiotic. Comparison of the structures of antibiotic-bound and antibiotic-free functional complexes enabled us to reveal structural rearrangements that take place in the PTC upon CHL binding as well as nascent peptide-induced changes in the drug binding site. Altogether, our structures provide the first view of CHL interactions not only with the ribosome but also with nascent protein chains and rationalize the observed context-specificity of action of this antibiotic. Importantly, our results also explain why other classes of PTC inhibitors, such as clinically relevant oxazolidinones, including the newest FDA-approved linezolid, while binding in the same site as CHL exhibit different context specificity of action.

## RESULTS AND DISCUSSION

### MTI/MAI tripeptide sequences can cause efficient CHL-dependent ribosome stalling

The main goal of this study is to gain understanding of the structural bases for the context specificity of CHL action, which may serve as a paradigm for other PTC-targeting drugs acting in a sequence-specific fashion, such as clinically important oxazolidinones. More specifically, we aimed to structurally decipher the role of the amino acid residue at the penultimate (−1) position of the growing polypeptide chain, which was shown to be critical for the action of CHL^17,18^. For our studies, we chose the short tripeptide CHL-stalling motif – Met-Thr-Ile (MTI) – which represents the three N-terminal amino acid residues of the *E. coli osmC* gene^17,19^. Biochemical and *in vivo* experiments suggested that, in the presence of CHL, the ribosome stalls when the third Ile codon in placed in the P site, likely because it fails to form a peptide bond between the donor substrate, peptidyl-tRNA carrying the MTI-tripeptide, and the incoming A-site acceptor substrate, histidyl-tRNA^17,19^. Using a solid-phase chemical synthesis of the short tRNA-substrate analogs mimicking the 3′-terminal CCA-end of the acceptor stem of full-length tRNA^20^, we generated the peptidyl-tRNA analog carrying the MTI-tripeptide (5’-ACCA-ITM) (**Figure S1**). In addition, because alanine residues are found more frequently than threonines (or serines) in the penultimate position of the nascent peptides associated with the sites of the most pronounced CHL action^17^, we also synthesized a peptidyl-tRNA analog carrying the MAI-tripeptide (5’-ACCA-IAM) (**Figure S1**). Lastly, as a negative control, we prepared an MFI-carrying peptidyl-tRNA analog because ribosome profiling analysis and biochemical experiments have shown that phenylalanine in the penultimate position of nascent chains does not support CHL-dependent ribosome stalling (**Figure S1**)^17,18^. Importantly, the peptidyl moieties of all three synthetic tRNA analogs were attached to the ribose of the A76-equivalent nucleotide via non-hydrolyzable amide bonds (**Figures 1A, B; Figure S1**), allowing us to capture the pre-reaction state in the structures.

**Figure 1.**
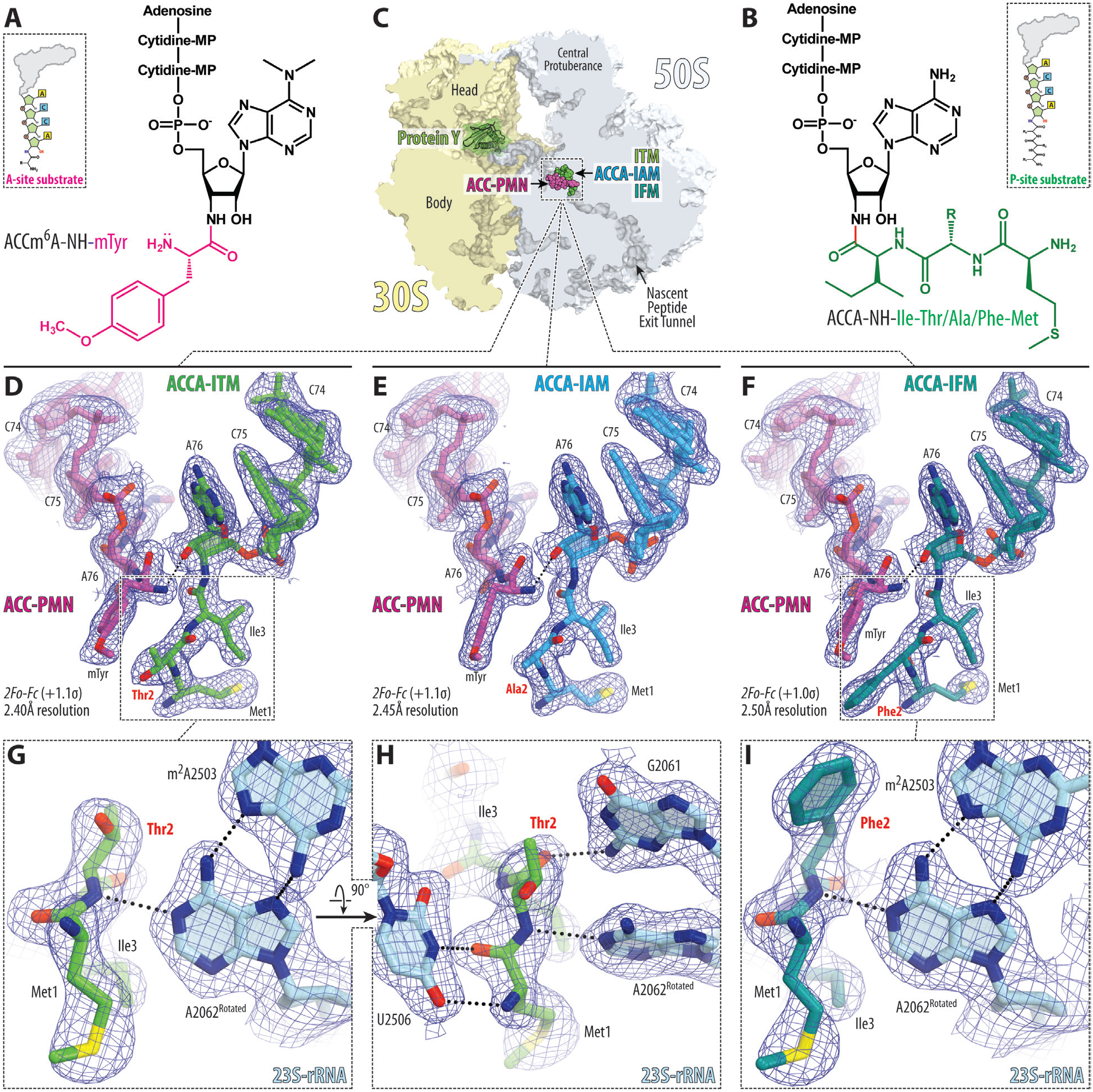
Electron density maps of the short tripeptidyl-tRNA analogs bound to the *T. thermophilus* 70S ribosome in the absence of CHL. (**A, B**) Chemical structures of the hydrolysis-resistant aminoacyl-tRNA mimic ACC-Puromycin (A), and the peptidyl-tRNA mimics ACCA-ITM/IAM/IFM (B). The amino acid moiety of ACC-Puromycin and peptide moiety of tripeptidyl-tRNA analogs are highlighted in purple and green, respectively. (**C**) Overview of the *T. thermophilus* 70S ribosome structures featuring short tRNA analogs viewed as a cross-cut section through the nascent peptide exit tunnel. The 30S subunit is shown in light yellow; the 50S subunit is in light blue; ribosome-bound protein Y is colored in green. (**D-F**) 2*F*_*o*_*-F*_*c*_ electron difference Fourier maps of ACC-PMN (purple) and either MTI-tripeptidyl-tRNA (D, green), or MAI-tripeptidyl-tRNA (E, blue), or MFI-tripeptidyl-tRNA (F, teal) analogs. The refined models of short tRNA analogs are displayed in their respective electron density maps after the refinement (blue mesh). The overall resolution of the corresponding structures and the contour levels of the depicted electron density maps are shown in the bottom left corner of each panel. (**G-I**) Close-up views of the MTI (G, H) or MFI (I) tripeptides interacting with the nucleotides of the 23S rRNA (light blue). Note that regardless of the nature of amino acid in the 2^nd^ position, all tripeptides have nearly identical paths in the ribosome exit tunnel and form a conserved network of H-bonds within the PTC.

Next, to ensure that the peptides included in the synthesized tRNA analogs indeed support (MTI and MAI) or counteract (MFI) CHL action during translation, we carried out primer extension inhibition assays (toe-printing), which allows detection of the drug-induced ribosome stalling site(s) along mRNAs with single-codon accuracy^19,21^. We used a template containing the first 27 codons of *osmC*, including the first three encoding the WT MTI sequence (*osmC*-MTI), as well as its two mutant versions encoding for the polypeptides starting with the MAI (*osmC*-MAI) or MFI (*osmC*-MFI) sequences. As expected, the addition of CHL to the cell-free translation system programmed with the *osmC* mRNA variants resulted in ribosome stalling at the Ile3 codon of the *osmC* ORF, when the threonine (*osmC*-MTI) or alanine (*osmC*-MAI) residues appeared in the penultimate position of the growing polypeptide chains, respectively (**Figure S2**, lanes 2 and 4, red arrowhead). In contrast, as expected, the replacement of Thr2 with Phe residue completely abolished CHL-dependent ribosome stalling (**Figure S2**, lane 6, red arrowhead), which is fully consistent with the reported context specificity of CHL action^17,18^. Thus, we utilized our synthetic MTI/MAI/MFI-tripeptidyl-tRNA analogs in our further structural studies.

### Structures of ribosome-bound tripeptidyl-tRNAs analogs reveal the uniform trajectories of their peptide chains

In order to provide a structural basis for the sequence-specific CHL-mediated ribosome stalling, we first explored whether the nature of the penultimate amino acid residue affects the trajectory of a nascent peptide within the 70S ribosome in the absence of antibiotic. To this end, we prepared a set of complexes containing ACC-puromycin (ACC-PMN) as the A-site substrate (**Figure 1A**) and one of the three tripeptidyl-tRNA analogs (**Figure 1B**) as the P-site substrate, crystallized them, and determined their structures (**Figure 1C**) at 2.4-2.5 Å resolution. To our knowledge, these structures provide the highest resolution of ribosome-donor substrate complexes reported to date. The observed excellent-quality electron density maps for both A- and P-site substrates in all three structures allowed the unambiguous modeling of the short aminoacyl- and peptidyl-tRNA analogs in the absence of antibiotic (**Figure 1D-I**). Superpositioning of all three structures revealed no significant differences in the overall conformations of the main peptide chain of the tripeptidyl tRNA analogs (**Figure 2A**). Not only the trajectories of the main-chain of the three tripeptides look the same, but even the positions of the main-chain and Cβ atoms of Thr2, Ala2, or Phe2 are identical (**Figure 2A**). Next, because we used short tRNA analogs, it was important to ensure that our new structures reflect functionally meaningful states. The alignment of our structures containing peptidyl-tRNA analogs with the previously published structures of the 70S ribosome in the pre-attack state containing either non-hydrolyzable amide-linked (**Figure 2B**)^22,23^ or native ester-linked full-length aa-tRNAs in the A- and P-sites (**Figure 2C**)^24^, shows essentially no structural differences in the position of the CCA moieties (**Figure S3A-C**) or the key 23S rRNA nucleotides around the PTC (**Figure S3D-F**). Thus, the position and orientation of the native ester bond linking the peptide to the 3’OH of the A76 residue of the P-site tRNA in the previous structures are nearly identical to the geometry of the amide linkages connecting the same moieties in our structures, suggesting that the short tripeptidyl-tRNA analogs fully mimic in structural terms the functionally-relevant pre-attack state of the peptidyl-tRNA. In particular, the orientation of the attacking α-amino group of the aa-tRNA or its analog (ACC-PMN) relative to the carbonyl carbon of the P-site substrate is indistinguishable between the structures harboring full-length tRNAs and our new structures with short tRNA analogs (**Figures 2B, C; S3A-C**). Moreover, the overall paths of the tripeptides of the peptidyl-tRNA analogs are similar to those of several other peptides in the nascent peptide exit tunnel (NPET), whose somewhat lower resolution structures were obtained previously using cryo-EM (**Figure S4C, D**)^25,26^.

**Figure 2.**
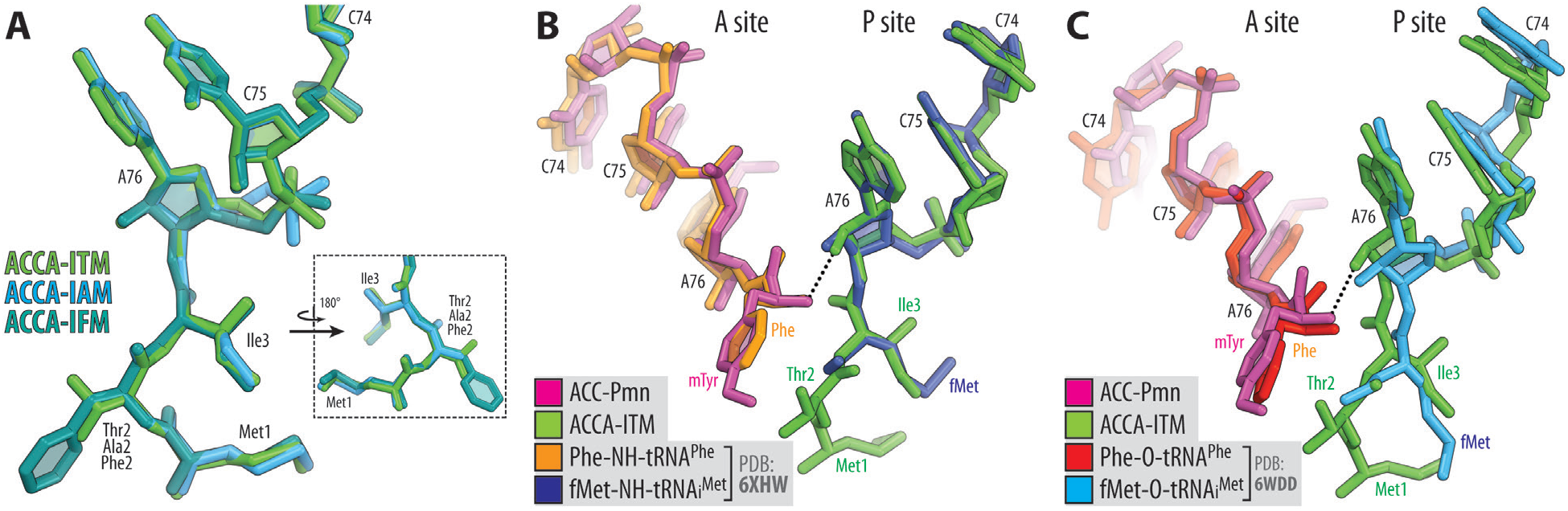
Comparison of the structures of short tripeptidyl-tRNA analogs reveals no significant differences between each other and with full-length tRNAs. (**A**) Superpositioning of the current 70S ribosome structures in complex with short tripeptidyl-tRNA analogs carrying MTI (green), MAI (blue), and MFI (teal) tripeptide sequences with each other. Note that the path of the growing polypeptide chain in the exit tunnel is not affected by the nature of the amino acid in the -1 position. (**B, C**) Comparison of our 70S ribosome structure carrying ACCA-PMN (magenta) and ACCA-ITM (green) short tRNA analogs in the A and P sites, respectively, with the previously reported structures of ribosome-bound full-length aminoacyl-tRNAs featuring either non-hydrolyzable amide linkages (B, PDB entry 6ZHW^23^) or native ester bonds (C, PDB entry 6WDD^24^) between the amino acid moiety and the ribose of A76 nucleotides. All structures were aligned based on domain V of the 23S rRNA. Note that, because the differences between the compared structures of the A- and P-site substrates are within experimental error, the short tRNA mimics represent functionally meaningful analogs of full-length tRNAs. Also, note that the amino acid moieties both in the A and P sites have very similar orientations regardless of whether or not the linkages are native or non-hydrolyzable.

The positions of the MTI/MAI/MFI peptide chains in the NPET are stabilized by several uniform hydrogen bonds (H-bonds) with the ribosome. For example, the exocyclic amino group of the G2061 residue of the 23S rRNA forms an H-bond with the main-chain carbonyl oxygen of the penultimate residue of the tripeptide (**Figure 1H**). We have also found that the presence of any of the three tripeptides (MTI, MAI, or MFI) causes the same re-orientation of the nucleotide A2062 of the 23S rRNA, which rotates by ∼160° relative to its placement in the vacant ribosome into a position where it forms a symmetric trans A-A Hoogsteen base pair with the residue A2503 (**Figure 1G, I**). In our structures, the main-chain amino group of the penultimate residue of the MTI/MAI/MFI-tripeptidyl moiety of the P-site substrate forms a clearly visible H-bond with the N1 atom of the A2062 residue, thereby stabilizing it in the rotated state (**Figure 1G-I**). Precisely the same rearrangement has been reported before to be caused by the ribosome-bound CHL antibiotic itself or its derivatives^10,27,28^. Our new structures provide the first experimental evidence that such rearrangement can be caused not necessarily by a drug molecule but also by a growing peptide chain irrespective of the nature of an amino acid in the penultimate position. These side-chain-independent and, therefore, likely uniform interactions of the main-chain groups of the growing peptide with the 23S rRNA nucleotides determine the observed orientation of the penultimate amino acid side chain towards the A site and potentially stabilize the transpeptidation-competent orientation of the terminal carbonyl group of the donor substrate during the peptidyl transfer reaction.

Altogether, the comparisons of the new structures revealed that the exact placement and the overall trajectories of all three tripeptides in the NPET of the 70S ribosome are strikingly similar (**Figure 2A**), irrespective of their ability to stall ribosomes in the presence of CHL (**Figure S2**). This critical result suggests that the observed context specificity of the CHL-induced ribosome stalling is not, due to alternative conformations of the nascent peptide chains within the NPET, as suggested previously^18^, neither it is due to differential peptide-induced conformational changes of the key nucleotides in the PTC acquired prior to the encounter with the drug (**Figure S3D-F**).

### Structures of ribosome-bound tripeptidyl-tRNAs and CHL rationalize its context specificity of action

The next step was to determine the structures of the same ribosomal complexes harboring short analogs of aminoacyl- and peptidyl-tRNAs, but now in the presence of CHL. However, after multiple attempts using high concentrations of the drug, we failed to detect any meaningful electron density for the ribosome-bound CHL in its canonical binding site^10^, while we observed clear electron density for the ACC-PMN in the A site and the MTI/MAI-tripeptidyl-tRNA analogs in the P site. The absence of CHL in the complexes is likely due to the clash between the methyl-tyrosine side chain of the ACC-PMN analog and the nitrobenzyl moiety of the CHL because they occupy exactly the same space in the A-site cleft of the ribosome, making their simultaneous presence on the ribosome impossible (**Figure S5A**). These data strongly suggest that the affinity of ACC-PMN for the ribosomal A site is high enough to outcompete CHL from its canonical binding site.

In order to avoid direct competition between the aminoacyl moiety of the A-site substrate ACC-PMN and the CHL molecule, we used the synthetic 5’-CACCA-3’ oligoribonucleotide, which lacks an attached amino acid and, therefore, mimics the 3’-end of a deacylated tRNA in the A site. Importantly, we found that the CACCA tRNA fragment stimulates the binding of the P-site tripeptidyl-tRNA analogs to the ribosome exactly the same way as ACC-PMN. Thus, using CACCA as a short A-site tRNA analog, we determined two structures of the ribosome in complex with CHL and either MTI or MAI stalling tripeptidyl-tRNA analogs at 2.50Å and 2.40Å resolution, respectively (**Figure 3A, B, D, E**). In both structures, the CHL molecule is bound to its canonical site in the PTC (**Figure S6**)^10^ so that the nitrobenzyl moiety of the drug intercalates into the hydrophobic pocket formed by the 23S rRNA residues A2451, C2452, and U2506 (**Figure S6D**). We were unable to obtain a similar structure with the non-stalling MFI-tripeptide, most likely because its bulky phenylalanine side chain competes with CHL for ribosome binding (**Figure S5B**). From the obtained structures, it became evident that the MTI or MAI stalling peptides could stabilize the ribosome-bound drug by directly interacting with it. With the MTI peptide, the hydroxyl group of the side chain of the penultimate threonine residue forms an H-bond with one of the two chlorine atoms of CHL (**Figure 3B, C**). Of note, the side chain of serine, which also conduces to CHL stalling when present in the penultimate position of a nascent chain^17^, would have the ability to form the same interaction with CHL. Similar to the threonine of MTI, the penultimate alanine of the MAI stalling peptide also stabilizes ribosome-bound CHL through an energetically favorable CH-π interaction between the side chain methyl group of alanine (serving as a CH-donor) and the CHL benzene ring (serving as a CH-acceptor) located 3.6Å away from each other (**Figure 3F**). The energy of CH-π type of interactions is estimated to be 1.5-2.5 kcal/mole^29^, which is comparable with the energy of standard H-bonds (0.5-1.8 kcal/mole)^30^, and should substantially increase the affinity of the drug to the ribosome. The observed specific contacts of the CHL molecule with the side chain of the penultimate residue of the MTI or MAI stalling peptides (**Figure 3C, F**) likely results in stronger binding of the drug to the ribosome, which presumably decreases its off-rate and makes it harder for the incoming aa-tRNA to displace the ribosome-bound antibiotic. Importantly, our structures reveal why it is specifically the penultimate amino acid residue (−1 position) of the peptidyl-tRNA that plays the defining role in CHL-induced ribosome stalling: the side chains of residues in position -2 (methionine) or position 0 (the isoleucine in the P site) protrude away from the CHL binding site, while it is only the side chain of residue at position -1 that faces the CHL binding pocket and the ribosomal A site (**Figure 3**). Therefore, our structural data rationalize previous ribosome profiling and biochemical findings^17^, showing that regardless of the nature of amino acids in other positions, it is the penultimate residue of the peptidyl-tRNA that can either interact or sterically interfere (**Figure S5B**) with the CHL in the PTC and, thus, affect its ability to stall ribosomes.

**Figure 3.**
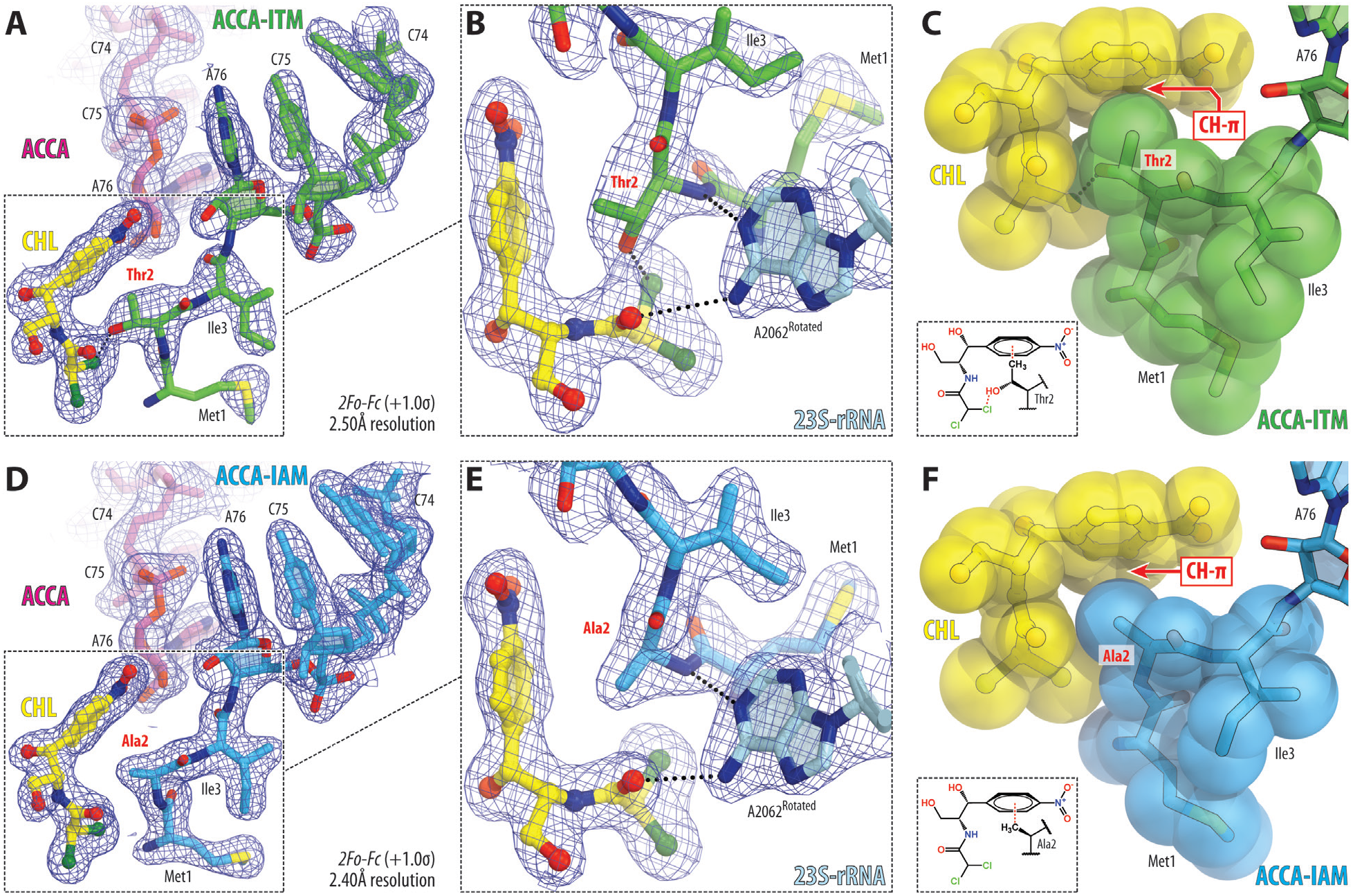
CHL directly interacts with the stalling peptides MTI and MAI in the *T. thermophilus* 70S ribosome. (**A, B, D, E**) 2*F*_*o*_*-F*_*c*_ electron density maps of the A-site ACCA (purple) and P-site short MTI-tripeptidyl-tRNA (A, B; green) or MAI-tripeptidyl-tRNA (A, B; blue) analogs bound to the *T. thermophilus* 70S ribosome in the presence of CHL (yellow). (**C, F**) Close-up views of the MTI (C) or MAI (F) tripeptides interacting with the CHL on the ribosome. Note that the side chains of penultimate threonine or alanine residues of the nascent peptide form H-bond or CH-π interactions with the dichloroacetic and/or nitrobenzyl moieties of CHL.

### Increased affinity model rationalizes CHL context-specificity of action

Our structural data allow us to unambiguously discriminate between the two recently proposed models – the increased affinity model and the drug-induced conformational change model^18^ – that were proposed to explain the observed context-specificity of CHL action *in vivo* and *in vitro*^17^. Our data support the *increased affinity model*. Apparently, the binding on-rate or affinity of CHL for the peptide-free ribosome is not high enough for the drug molecule to significantly compete with an incoming aminoacyl-tRNA, which is reflected in poor drug-induced ribosome stalling during the first few rounds of elongation on many mRNA templates^17,19^. However, once the emerging peptide chain reaches a certain length (3-6 amino acids), it becomes better anchored in the NPET and could now provide an additional binding interface for the CHL molecule, especially if the chain carries alanine, serine, or threonine in the penultimate position, which serves as specific interacting partners for CHL, increasing its affinity for the ribosome and possibly stimulating the rate of drug binding. The CHL molecule, now firmly anchored in the A site due to its interaction with the penultimate residue of the stalling peptide, would prevent accommodation of the aminoacyl moiety of the aa-tRNA into the A-site cleft, thereby inhibiting peptidyl transfer (**Figure 4A**). As revealed by smFRET studies, the ribosome with tightly bound CHL can fully accommodate the body of the incoming aa-tRNA but is unable to place the acceptor amino acid in the conformation required for productive peptide bond formation, resulting in the tRNA rejection and continued rounds of aminoacyl-tRNA sampling (**Figure 4A**) (Choi et al., 2019). Based on our structural data, residues with small side chains such as Ala, Ser, or Thr in the penultimate position of the growing peptide chain are conducive to CHL-dependent ribosome stalling not only because their side chains do not interfere with CHL binding as those of large amino acids might do, but also because their side chains additionally facilitate CHL binding by establishing specific interactions with the drug. In contrast, residues with larger side chains in the penultimate position of the growing peptide are sterically incompatible with CHL and either prevent its binding to the ribosome or cause its dissociation (**Figure 4B**). This steric incompatibility of the ribosome-bound CHL and the bulky side chains at the penultimate position of the nascent peptide explains the previously mysterious inability of CHL to fully inhibit puromycin reaction on polysomes^14^. Indeed, at any given moment, at least some of the translating ribosomes of the polysome fraction should contain growing peptides with bulky side chains at the penultimate position preventing CHL binding and hence, inhibition. At the same time, none of these bulky residues interfere with the binding and action of puromycin or an incoming aa-tRNA. Similarly, ribosome profiling studies have shown that adding chloramphenicol to bacterial cells prior to cell lysis still allows the movement of ribosomes by a few codons^17,31,32^, potentially reflecting the poor ability of the drug to bind to the ribosome carrying the unfavorable nascent chains.

**Figure 4.**
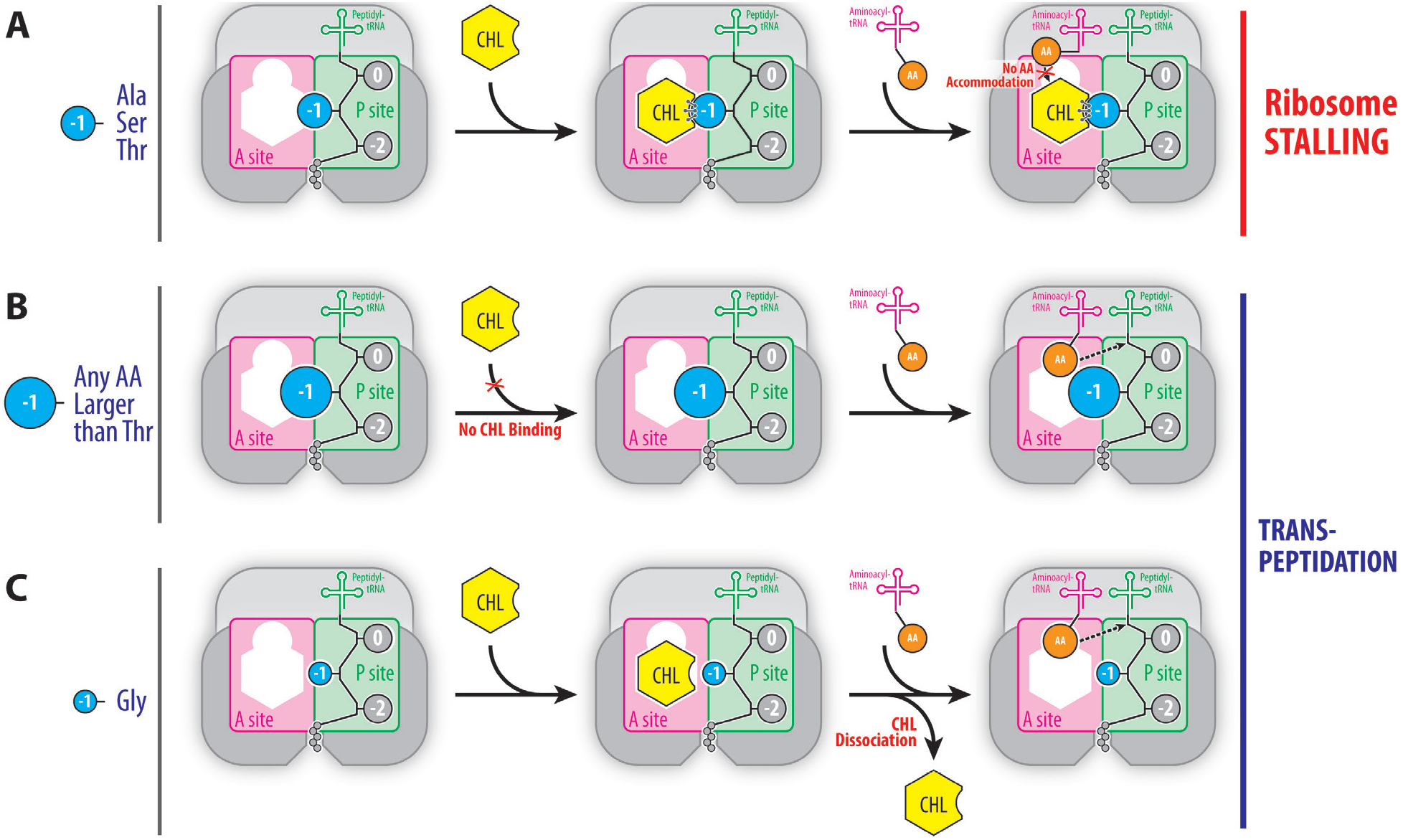
Schematic diagram illustrating the *increased affinity model*. (**A**) When the growing polypeptide chain carries alanine, serine, or threonine (blue circle) in the penultimate position, the affinity of CHL (yellow hexagon) for the ribosome increases due to direct interactions with the side chains of these amino acids preventing accommodation of the aminoacyl moiety of the aa-tRNA into the A site and, thereby, inhibiting peptidyl transferase reaction, which results in ribosome stalling. (**B**) Amino acid residues with larger side chains in the penultimate position of the growing peptide are sterically incompatible with CHL preventing its binding to the ribosome. In the absence of ribosome-bound CHL, aa-tRNA normally accommodates into the ribosomal A site resulting in an unperturbed transpeptidation reaction. (**C**) Glycine residue in the penultimate position of the growing peptide does not interfere with the binding of CHL but also does not increase the drug’s affinity for the ribosome. As a result, loosely bound CHL is displaced from the ribosomal A site by the incoming aa-tRNA, which, after accommodation, can normally react with the P-site substrate. Small and large ribosomal subunits are highlighted in shades of grey. A and P sites are highlighted in magenta and green, respectively. Amino acids of the growing polypeptide chain are shown by grey circles with the penultimate amino acid residue highlighted in blue.

Apparently, glycine residue in the penultimate position of the growing peptide would not interfere with CHL binding due to the lack of a side chain (**Figure S7A**) but, at the same time, would not be able to establish an H-bond or a CH-π interaction with the CHL molecule and, hence, is unlikely to increase drug’s affinity for the ribosome (**Figure 4C**). Nevertheless, with a lesser frequency compared to Ala/Ser/Thr, glycines do appear at the penultimate position of the CHL-dependent ribosome stalling sites revealed by ribosome profiling experiments^17^ as well as by our toe-printing analysis (**Figure S2**, lanes 2, 4, and 6, green arrowhead), suggesting that absence of interference between the growing peptide and the drug is the primary feature of CHL-stalling motifs. In other words, even if glycine in the penultimate position of the growing peptide chain does not increase the affinity of CHL for the ribosome, at least it does not interfere with CHL binding, that is in contrast to all remaining amino acid residues whose side chains are bulkier than those of Ser or Thr. For example, cysteines and valines have the smallest side chains out of all remaining amino acids and might seem to be comparable in size with Ser or Thr, respectively. However, *in silico* modeling shows that, due to the 0.4 Å-larger van-der-Waals radius of a sulfur atom compared to oxygen, the cysteine residue is actually larger than serine or threonine and can barely co-exist with the ribosome-bound CHL molecule (**Figure S7B**). *In silico* modeling of valine shows that one of its terminal methyl groups clash with the dichloroacetic moiety of CHL, while the other methyl group comes too close to the CHL benzene ring (**Figure S7C**). Altogether, our structural analysis suggests that cysteine, valine, or any other residue with a larger side chain in the penultimate position of the growing peptide cannot co-exist with the ribosome-bound CHL and, thus, prevents its binding to the ribosome.

Remarkably, while CHL is one of the oldest known PTC-targeting antibiotics and is often viewed as a prototype of this class, members of the newest classes of clinically important PTC-targeting protein synthesis inhibitors that are chemically unrelated to CHL, such as linezolid (LZD), bind and act upon the PTC of the bacterial ribosome in a very similar fashion (**Figure S8A**)^17,18,33^. Curiously, LZD exhibits a pronounced context-specific mode of action similar to that of CHL: both drugs effectively inhibit peptide bond formation one step after an alanine residue appears in the growing polypeptide chain^17,18^. However, the main difference in the context specificity between the two drugs is that, unlike CHL, LZD does not cause ribosome stalling when serine or threonine appears in the penultimate position of the nascent chain^17,18^. To rationalize the observed strong preference of LZD for alanines, we superimposed the previous structure of ribosome-bound LZD^33^ with the new structures of MAI- or MTI-tripeptidyl-tRNA analogs (**Figure S8B, C**). Interestingly, the aromatic benzene rings of these two chemically distinct drug molecules superimpose well with each other (**Figure S8A**), suggesting that the side chain of alanine in the penultimate position of the growing polypeptide chain likely forms the same CH-π interactions with LZD as it does with CHL (**Figure S8B**), thereby increasing its affinity for the ribosome. At the same time, any side chains larger than alanine, including serine and threonine, likely clash either with the acetyl-amino-methyl tail or the fluorobenzene ring of LZD (**Figure S8C**), making drug binding to such ribosome nascent chain complex highly unfavorable. In summary, our *in silico* analysis reveals that only the alanine side chain in the penultimate position of the growing peptide can interact with the ribosome-bound LZD, while all other bulkier side chains are incompatible with the drug.

## CONCLUSIONS

In contrast to the previous structural studies that utilized the vacant ribosome, our functional complexes show that true the drug binding site is actually formed not only by the ribosome alone but also by the growing polypeptide chain, and the shape of the drug-binding pocket continuously changes (which, in turn, impacts the antibiotic affinity) as the nascent chain is synthesized by the ribosome. Our results demonstrate that the interplay between the ribosome, the nascent peptide chain, and the ribosome-bound drug defines whether or not translation stalling would occur in the presence of the PTC-targeting antibiotics such as CHL or LZD. The new structures reported here rationalize the previous data showing that the placement of alanine, serine, or threonine in the penultimate position of the nascent peptide chain is required for the CHL-induced ribosome stalling^17,18^. Our structural analysis reveals that the presence of a non-bulky side chain in the penultimate position of a growing peptide allows unhindered access of the drug to its canonical binding site. Moreover, the side chains of such amino acids as alanine, threonine, and likely serine in this position directly interact with the ribosome-bound CHL, further increasing its affinity to the ribosome. In our view, the CHL-induced ribosome stalling occurs because the aminoacyl moiety of an incoming A-site tRNA is unable to displace the tightly bound CHL molecule from its canonical binding site, which happens only when it is stabilized by interactions with the proper penultimate residue of the growing peptide, and, therefore, cannot get accommodated into the ribosomal A site making peptide bond formation impossible^18^. From this perspective, the nascent peptide chain plays the role of drug affinity modulator^18^ and should be considered as a part of the CHL or LZD binding site. Thus, these drugs should no longer be considered as competitive ribosome inhibitors but rather classified as uncompetitive inhibitors – a rare class of enzyme inhibitors, which bind adjacent to the active site of an enzyme and require a substrate to be present prior to their binding (e.g., inhibition of inosine-monophosphate dehydrogenase by mycophenolic acid^34^). In summary, the structures reported here provide the first insight into the mechanism of context-specificity of drug action and also provide a previously unmatched level of structural detalization of nascent peptide chains, revealing that their paths in the NPET are uniform despite their unique sequences.

## AUTHOR CONTRIBUTIONS

L.F. and R.M. synthesized short tripeptidyl-tRNA mimics; D.K. and N.V.-L. performed the toe-printing analysis; E.A.S. and Y.S.P designed and performed X-ray crystallography experiments; R.M., and Y.S.P. supervised the experiments. All authors interpreted the results. E.A.S., R.M. and Y.S.P. wrote the manuscript.

## ACKNOWLEDGMENTS

We thank Julia Thaler (Innsbruck) for synthetic support on the RNA-peptide conjugates. We thank Dr. Alexander Mankin for critical reading of the manuscript, as well as Dr. Kiira Ratia and Dr. Maxim Svetlov for valuable suggestions. We thank the staff at NE-CAT beamlines 24ID-C and 24ID-E for help with data collection and freezing of the crystals, especially Drs. Malcolm Capel, Frank Murphy, Igor Kourinov, Anthony Lynch, Surajit Banerjee, David Neau, Jonathan Schuermann, Narayanasami Sukumar, James Withrow, Kay Perry, and Cyndi Salbego.

This work is based upon research conducted at the Northeastern Collaborative Access Team beamlines, which are funded by the National Institute of General Medical Sciences from the National Institutes of Health [P41 GM103403 to NE-CAT]. The Pilatus 6M detector on 24-ID-C beamline is funded by a NIH-ORIP HEI [S10-RR029205 to NE-CAT]. The Eiger 16M detector on 24-ID-E beamline is funded by a NIH-ORIP HEI grant [S10-OD021527 to NE-CAT]. This research used resources of the Advanced Photon Source, a U.S. Department of Energy (DOE) Office of Science User Facility operated for the DOE Office of Science by Argonne National Laboratory under Contract No. DE-AC02-06CH11357.

This work was supported by Illinois State startup funds [to Y.S.P.], National Institutes of Health [R01-GM132302 to Y.S.P.], National Science Foundation [MCB-1951405 to N.V.-L.], and Austrian Science Fund FWF [P31691 and F8011 to R.M.].

## ACCESSION NUMBERS

Coordinates and structure factors were deposited in the RCSB Protein Data Bank with accession codes:

- **7XXX** for the *T. thermophilus* 70S ribosome in complex with protein Y, A-site aminoacyl-tRNA analog ACC-Pmn, and P-site peptidyl-tRNA analog **ACCA-ITM**;
- **7YYY** for the *T. thermophilus* 70S ribosome in complex with protein Y, A-site aminoacyl-tRNA analog ACC-Pmn, and P-site peptidyl-tRNA analog **ACCA-IAM**;
- **7ZZZ** for the *T. thermophilus* 70S ribosome in complex with protein Y, A-site aminoacyl-tRNA analog ACC-Pmn, and P-site peptidyl-tRNA analog **ACCA-IFM**;
- **7UUU** for the *T. thermophilus* 70S ribosome in complex with protein Y, A-site deacylated tRNA analog CACCA, P-site peptidyl-tRNA analog **ACCA-ITM**, and **chloramphenicol**;
- **7VVV** for the *T. thermophilus* 70S ribosome in complex with protein Y, A-site deacylated tRNA analog CACCA, P-site peptidyl-tRNA analog **ACCA-IAM**, and **chloramphenicol**;

## REFERENCES

1. Blanchard, S.C., Cooperman, B.S. & Wilson, D.N. Probing translation with small-molecule inhibitors. Chem. Biol. 17, 633–645 (2010).

2. Barnhill, A.E., Brewer, M.T. & Carlson, S.A. Adverse effects of antimicrobials via predictable or idiosyncratic inhibition of host mitochondrial components. Antimicrob. Agents Chemother. 56, 4046–4051 (2012).

3. Jones, C.N., Miller, C., Tenenbaum, A., Spremulli, L.L. & Saada, A. Antibiotic effects on mitochondrial translation and in patients with mitochondrial translational defects. Mitochondrion 9, 429–437 (2009).

4. Singh, R., Sripada, L. & Singh, R. Side effects of antibiotics during bacterial infection: mitochondria, the main target in host cell. Mitochondrion 16, 50–54 (2014).

5. Li, C.H., Cheng, Y.W., Liao, P.L., Yang, Y.T. & Kang, J.J. Chloramphenicol causes mitochondrial stress, decreases ATP biosynthesis, induces matrix metalloproteinase-13 expression, and solid-tumor cell invasion. Toxicol. Sci. 116, 140–150 (2010).

6. Cohen, B.H. & Saneto, R.P. Mitochondrial translational inhibitors in the pharmacopeia. Biochim. Biophys. Acta 1819, 1067–1074 (2012).

7. Syriopoulou, V.P., Harding, A.L., Goldmann, D.A. & Smith, A.L. In vitro antibacterial activity of fluorinated analogs of chloramphenicol and thiamphenicol. Antimicrob. Agents Chemother. 19, 294–297 (1981).

8. Dunkle, J.A., Xiong, L., Mankin, A.S. & Cate, J.H. Structures of the Escherichia coli ribosome with antibiotics bound near the peptidyl transferase center explain spectra of drug action. Proc. Natl. Acad. Sci. USA 107, 17152–17157 (2010).

9. Bulkley, D., Innis, C.A., Blaha, G. & Steitz, T.A. Revisiting the structures of several antibiotics bound to the bacterial ribosome. Proc. Natl. Acad. Sci. USA 107, 17158–17163 (2010).

10. Svetlov, M.S. et al. High-resolution crystal structures of ribosome-bound chloramphenicol and erythromycin provide the ultimate basis for their competition. RNA 25, 600–606 (2019).

11. Drainas, D., Kalpaxis, D.L. & Coutsogeorgopoulos, C. Inhibition of ribosomal peptidyltransferase by chloramphenicol. Kinetic studies. Eur. J. Biochem. 164, 53–58 (1987).

12. Kucan, Z. & Lipmann, F. Differences in chloramphenicol sensitivity of cell-free amino acid polymerization systems. J. Biol. Chem. 239, 516–520 (1964).

13. Vazquez, D. Antibiotics affecting chloramphenicol uptake by bacteria. Their effect on amino acid incorporation in a cell-free system. Biochim. Biophys. Acta 114, 289–295 (1966).

14. Cannon, M. The puromycin reaction and its inhibition by chloramphenicol. Eur. J. Biochem. 7, 137–145 (1968).

15. Lovett, P.S. Translation attenuation regulation of chloramphenicol resistance in bacteria -a review. Gene 179, 157–162 (1996).

16. Lovett, P.S. Translational attenuation as the regulator of inducible cat genes. J. Bacteriol. 172, 1–6 (1990).

17. Marks, J. et al. Context-specific inhibition of translation by ribosomal antibiotics targeting the peptidyl transferase center. Proc. Natl. Acad. Sci. USA 113, 12150–12155 (2016).

18. Choi, J. et al. Dynamics of the context-specific translation arrest by chloramphenicol and linezolid. Nat. Chem. Biol. 16, 310–317 (2020).

19. Orelle, C. et al. Tools for characterizing bacterial protein synthesis inhibitors. Antimicrob. Agents Chemother. 57, 5994–6004 (2013).

20. Moroder, H. et al. Non-hydrolyzable RNA-peptide conjugates: a powerful advance in the synthesis of mimics for 3’-peptidyl tRNA termini. Angew. Chem. Int. Ed. Engl. 48, 4056–4060 (2009).

21. Hartz, D., McPheeters, D.S., Traut, R. & Gold, L. Extension inhibition analysis of translation initiation complexes. Methods Enzymol. 164, 419–425 (1988).

22. Polikanov, Y.S., Steitz, T.A. & Innis, C.A. A proton wire to couple aminoacyl-tRNA accommodation and peptide-bond formation on the ribosome. Nat. Struct. Mol. Biol. 21, 787–793 (2014).

23. Svetlov, M.S. et al. Structure of Erm-modified 70S ribosome reveals the mechanism of macrolide resistance. Nat. Chem. Biol. 17, 412–420 (2021).

24. Loveland, A.B., Demo, G. & Korostelev, A.A. Cryo-EM of elongating ribosome with EF-Tu*GTP elucidates tRNA proofreading. Nature 584, 640–645 (2020).

25. Herrero Del Valle, A. et al. Ornithine capture by a translating ribosome controls bacterial polyamine synthesis. Nat. Microbiol. 5, 554–561 (2020).

26. Su, T. et al. The force-sensing peptide VemP employs extreme compaction and secondary structure formation to induce ribosomal stalling. Elife 6(2017).

27. Tereshchenkov, A.G. et al. Binding and action of amino acid analogs of chloramphenicol upon the bacterial ribosome. J. Mol. Biol. 430, 842–852 (2018).

28. Chen, C.W. et al. Binding and action of triphenylphosphonium analog of chloramphenicol upon the bacterial ribosome. Antibiotics (Basel) 10(2021).

29. Nishio, M. The CH/pi hydrogen bond in chemistry. Conformation, supramolecules, optical resolution and interactions involving carbohydrates. Phys. Chem. Chem. Phys. 13, 13873–13900 (2011).

30. Fersht, A.R. The hydrogen bond in molecular recognition. Trends in Biochemical Sciences 12, 301–304 (1987).

31. Becker, A.H., Oh, E., Weissman, J.S., Kramer, G. & Bukau, B. Selective ribosome profiling as a tool for studying the interaction of chaperones and targeting factors with nascent polypeptide chains and ribosomes. Nat. Protoc. 8, 2212–2239 (2013).

32. Mohammad, F., Woolstenhulme, C.J., Green, R. & Buskirk, A.R. Clarifying the translational pausing landscape in bacteria by ribosome profiling. Cell Rep. 14, 686–694 (2016).

33. Ippolito, J.A. et al. Crystal structure of the oxazolidinone antibiotic linezolid bound to the 50S ribosomal subunit. J. Med. Chem. 51, 3353–3356 (2008).

34. Hedstrom, L. IMP dehydrogenase: structure, mechanism, and inhibition. Chem. Rev. 109, 2903–2928 (2009).

