## Supplementary Information for "Structural basis for the context-specific action of classic peptidyl transferase inhibitors"

**This file includes:**

I. Materials and Methods;
II. Supplementary Table 1;
III. Supplementary Figures 1 to 7 with legends;
IV. Supplementary References.

### I. MATERIALS AND METHODS

***Synthesis of hydrolysis-resistant A-site aminoacyl- and deacyl-tRNA analogs.***

Synthetic aminoacyl-tRNA mimic, adenosine-cytidine-cytidine-puromycin (ACC-PMN) was obtained from Horizon Discovery Inc. (Chicago, USA). Synthetic deacyl-tRNA analog, pentanucleotide 5'-CACCA-3', was obtained from Integrated DNA Technologies (Coralville, IA).

***Synthesis of hydrolysis-resistant peptidyl-tRNA analogs.***

The tripeptidyl-tRNA conjugates featuring MTI, MAI, ot MFI peptide sequences were produced as outlined in **Figure S1A** and as described below. The DMTO-rA3′-NH-Ile-NHFmoc solid support **1** (**Figure S1A**) for the synthesis of ACCA-Ile-Ala-Met, ACCA-Ile-Thr-Met, and ACCA-Ile-Phe-Met conjugates was produced according to the following references[1-3]. The assembly of the conjugates was based on Fmoc peptide solid-phase synthesis and RNA solid-phase synthesis using 2'-*O*-[(triisopropylsilyl)oxy]methyl (TOM) protected nucleoside building blocks as outlined in **Figure S1A** and as described below, essentially based on the following references[4]:

1. ***Solid-phase peptide assembly on the solid support 1***. In an ABI synthesis cartridge, solid support **1** (40 mg) was soaked with dry *N*,*N*-dimethylformamide (2 mL, 30 min). For deprotection of the *N*α-Fmoc group, the solid support was treated twice with piperidine solution (20 % in *N*,*N*-dimethylformamide, 1.5 mL, 2 min, 15 min) and subsequently washed with *N*,*N*-dimethyl-formamide (3 x 2 mL). Coupling was performed by adding *N*,*N*-diisopropylethylamine (17 μL ) to a solution (500 μL) of 100 mM (1-cyano-2-ethoxy-2-oxoethylidenaminooxy)dimethylamino-morpholino-carbenium-hexafluorophosphate and Fmoc-protected amino acid (100 μmol) in *N*,*N*-dimethylformamide. After 1 minute of preactivation, the solid support was treated with this solution for 1.5 hours. This step was performed twice. Then, the solid support was washed with *N*,*N*-dimethylformamide (3 x 2 mL), methanol (3 x 2 mL), and dichloromethane (3 x 2 mL) and dried on high vacuum.
2. ***Solid-phase RNA assembly on the tripeptidyl solid support 1.*** The ACCA moiety was assembled using a *Nucleic Acid Synthesizer* (*ABI 392)* following standard synthesis protocols. Detritylation (120 s): dichloroacetic acid/1,2-dichloroethane (4/96); coupling (360 s): phosphoramidites (0.1 M in acetonitrile) were activated with benzylthiotetrazole (0.3 M in acetonitrile); capping (2 x 10 s, Cap A/Cap B = 1/1): Cap A: 0.49 M *N,N*-dimethylaminopyridine in acetonitrile, Cap B: acetic anhydride, 2,4,6-collidine, acetonitrile (10/15/25); oxidation (20 s): $I_2$ (0.2 M) in tetrahydrofuran/pyridine/$H_2O$

(35/10/5). Amidites, benzylthiotetrazole, and capping solutions were dried over activated molecular sieves (4Å) overnight.

3. ***Deprotection of the 3'-tripeptidyl-ACCA conjugates***. (**A**) *Fmoc and cyanoethyl deprotection*. In the ABI synthesis column, the solid support was treated with a solution of 20 % piperidine in acetonitrile (5 mL, 5 min), washed with acetonitrile (3 x 2 mL), and dried. (**B**) *Acyl deprotection and cleavage from the solid support*. For the conjugates synthesized on solid support **1**, the beads were transferred into a screwcap vial, and equal volumes of aqueous methylamine (33% m/m), and concentrated aqueous ammonia (28-30% m/m) were added. After 20 minutes at 65 °C, the supernatant was filtered and evaporated to dryness. (**C**) *2'-O-TOM deprotection*. The obtained residue was treated with tetrabutylammonium fluoride trihydrate ($TBAF \cdot 3H_2O$) in tetrahydrofuran (1 M, 1 mL) overnight at room temperature. The reaction was quenched by the addition of triethylammonium acetate (TEAA) (1 M, pH 7.4, 1 mL). After evaporating tetrahydrofuran, the solution was applied on a size-exclusion chromatography column (*GE Healthcare*, HiPrep 26/10 Desalting, 2.6 x 10 cm, Sephadex G25). By eluting with $H_2O$, the conjugate-containing fractions were collected, evaporated to dryness, and the residue was dissolved in $H_2O$ (1 mL). Analysis of the crude products was performed by anion-exchange chromatography on a Dionex DNAPac PA-100 column (4 x 250 mm) at 80°C. Flow rate: 1 mL $min^{-1}$; eluent A: 25 mM Tris-HCl (pH 8.0) and 20 mM $NaClO_4$ in 20% aqueous acetonitrile, eluent B: 25 mM Tris-HCl (pH 8.0) and 0.60 M $NaClO_4$ in 20% aqueous acetonitrile; gradient: 0-35 % B in A within 28 min; UV detection at $\lambda = 260$ nm.
4. ***Purification of the 3'-tripeptidyl-ACCA conjugate.*** The crude deprotected conjugates were purified on a semipreparative Dionex DNAPac PA-100 column (9 x 250 mm) at 80 °C with a flow rate of 2 mL $min^{-1}$ (for eluents see above). Fractions containing the conjugate were concentrated to near dryness and diluted with 0.1 M $(Et_3NH)^+HCO_3^-$ and loaded on a C18 SepPak Plus cartridge (Waters/Millipore), washed with $H_2O$, and eluted with $H_2O/CH_3CN$ (1:1). Conjugate-containing fractions were evaporated to dryness and dissolved in $H_2O$ (0.5 mL). The quality of the purified conjugate was analyzed by analytical anion-exchange chromatography (for conditions, see above). The molecular weight of the synthesized conjugate was confirmed by LC-ESI mass spectrometry (**Figure S1B**). Yields were determined by UV photometrical analysis of conjugate solutions.

***Toe-printing analysis.***

For the toe-printing analysis of the CHL-mediated ribosome stalling during *in vitro* translation at the third codon of the *osmC* open reading frame[5], we used the following DNA template that contains the T7 promoter region, the original first 27 codons (underlined) of the *osmC* gene, followed by a stop codon (red), and an annealing region for the toe-printing primer NV1:

> ATTAATACGACTCACTATAGGGATATAAGGAGGAAAACAT ATG **ACA** ATC CAT AAG AAA GGT CAG GCA CAC TGG GAA GGC GAT ATC AAA CGC GGG AAG GGA ACA GTA TCC ACC GAG AGT GGC TAA GCTCTTTGGTTAATAAGCAAAATTCATTATAACC

This DNA template was prepared by PCR using T7-ermCL-osmC-fwd (5'-ATTAATACGACTCACTATAGGGATATAAGGAGGAAAACATATGACAATCCATAAGAAAGG) and osmC-NV1-reverse (5'-GGTTATAATGAATTTTGCTTATTAACCAAAGAGCTTAGCCACTCTCGGTGGATACTGTTC) primers. DNA templates containing mutations of the 2nd Thr codon (ACA, highlighted in bold in the above sequence) to either alanine (GCA) or phenylalanine (TTC) were prepared by combining forward primer T7-ermCL-osmC-T2A-fwd (5'-ATTAATACGACTCACTATAGGGATATAAGGAGGAAAACATATGGCAATCCATAAGAAAGG) or T7-ermCL-osmC-T2F-fwd (5'-ATTAATACGACTCACTATAGGGATATAAGGAGGAAAACATATGTTCATCCATAAGAAAGG), respectively, with the same osmC-NV1-reverse primer. Toe-printing analysis was performed essentially as described before[6]. Translation time was 5 min and reactions were supplemented with additional 3 mM $MgCl_2$. When added, CHL was at 200 μM final concentration.

***Crystallographic structure determination.***

To obtain high-resolution structures that would allow visualizing of structural features crucial for this study, we took advantage of some of the recent findings by our group. In our initial trials, we tried to either co-crystallize the tripeptidyl-tRNA analogs together with the 70S ribosome or to soak them into the pre-formed ribosome crystals hoping that the analogs would bind into the ribosomal P site leaving the A site vacant. Unfortunately, these initial attempts yielded only poor electron density corresponding to any of the three synthetic P-site substrates tested. Increased resolution and significantly better electron density maps resulted from the addition of the aminoacyl-tRNA analog 5'-ACC-puromycin (ACC-PMN) (**Figure 1A**), which we have shown previously that strongly binds to the A site of the ribosome and fully mimicks aminoacyl-tRNA[7,8]. Further improvement came from the use of the protein Y (PY), which binds to the small ribosomal subunit and is known to improve the resolution of the ribosome complexes[9-12]. A specific advantage of using the PY in this study results from its ability to compete with the binding of intact tRNAs to the ribosome, which helped to purge any residual ribosome-bound tRNAs that were carried over during the ribosome purification. In the absence of the competing full-length tRNAs, short peptidyl- and aminoacyl-tRNA analogs freely bind to the A and P sites on the large ribosomal subunit and can be visualized using X-ray crystallography.

Complexes of the wild-type *Tth* 70S ribosomes with PY, and short hydrolysis-resistant tRNA analogs were formed as described previously[7-10]. The short aminoacyl-tRNA analog ACC-PMN and deacylated tRNA analog ACCA were added to 100 μM final concentration, which corresponds to 20-fold molar excess over the 70S ribosomes. The short tripeptidyl-tRNA analogs were added to 50 μM final concentration. For *Tth* 70S ribosome complexes with CHL, the drug was included in the crystallization mixture (500 μM) and then later added to the stabilization buffers. Crystallization, crystal stabilization, and freezing as well as collection and processing of the X-ray diffraction data, model building, and structure refinement were performed as described in our previous publications[8,10,13-15]. Structural model and restraints for CHL were generated using PRODRG online software (http://prodrg1.dyndns.org)[16]. The statistics of data collection and refinement are compiled in **Table S1**. All figures showing atomic models were generated using the PyMol software (www.pymol.org).

Importantly, unlike all previous ribosome structures harboring *in cis* synthesized peptidyl-tRNAs (for example[17,18]), the peptides in our complexes were not made by the very ribosomes that were crystallized and, instead, were introduced *in trans*. Although previously it has been already demonstrated that peptidyl-tRNAs can be introduced *in trans* and sustain protein synthesis[19], here we provide the first evidence that the structures of the peptide backbones of the *in cis* synthesized and *in trans* introduced peptidyl-tRNAs are nearly identical and, therefore, such in-trans-introduced peptidyl-tRNAs represent a functionally meaningful state (**Figure S4C, D**). Therefore, the new ribosome structures harboring short aminoacyl- and peptidyl-tRNA analogs adequately reflect the pre-attack state of the peptidyl transferase center and can be used as a model for stalling complexes.

#### II. SUPPLEMENTARY TABLES

##### Table S1. X-ray data collection and refinement statistics.

| ***Crystals*** | | **70S-PY complex with ACCA-ITM and ACC-PMN** | **70S-PY complex with ACCA-IAM and ACC-PMN** | **70S-PY complex with ACCA-IFM and ACC-PMN** |
|---|---|---|---|---|
| ***Diffraction data*** | | | | |
| Space Group | | $P2_12_12_1$ | $P2_12_12_1$ | $P2_12_12_1$ |
| Unit Cell Dimensions, Å (a x b x c) | | 209.64 x 448.67 x 619.79 | 209.61 x 448.45 x 618.45 | 209.32 x 449.63 x 620.21 |
| Wavelength, Å | | 0.9795 | 0.9795 | 0.9795 |
| Resolution range (outer shell), Å | | 224-2.40 (2.46-2.40) | 309-2.45 (2.51-2.45) | 310-2.50 (2.56-2.50) |
| I/σI (outer shell) | | 7.05 (0.92) | 6.75 (1.00) | 9.53 (1.07) |
| Resolution at which I/σI=1, Å | | 2.40 | 2.45 | 2.50 |
| Resolution at which I/σI=2, Å | | 2.65 | 2.67 | 2.70 |
| CC(1/2) at which I/σI=1, % | | 14.4 | 21.8 | 17.4 |
| CC(1/2) at which I/σI=2, % | | 48.3 | 51.6 | 44.4 |
| Completeness (outer shell), % | | 99.7 (98.7) | 99.9 (100.0) | 99.9 (99.7) |
| $R_{merge}$ (outer shell)% | | 18.3 (147.4) | 20.7 (208.0) | 24.8 (240.9) |
| No. of crystals used | | 1 | 1 | 2 |
| No. of Reflections Used: | Observed | 12,556,809 | 14,684,473 | 23,964,58, |
| | Unique | 2,234,911 | 2,100,555 | 1,986,436 |
| Redundancy (outer shell) | | 5.62 (5.03) | 6.99 (7.20) | 12.06 (11.89) |
| ***Refinement*** | | | | |
| Resolution range of the diffraction data included in the refinement, Å | | 182-2.40 | 187-2.45 | 155-2.50 |
| $R_{work}/R_{free}$, % | | 23.4/29.1 | 20.0/24.7 | 22.7/28.1 |
| *No. of Non-Hydrogen Atoms* | | | | |
| RNA | | 192,629 | 192,629 | 192,629 |
| Protein | | 93,149 | 93,145 | 93,157 |
| Ions (Mg, K, Zn, Fe) | | 2,551 | 2,551 | 2,551 |
| Waters | | 9,506 | 9,504 | 9,497 |
| *Ramachandran Plot* | | | | |
| Favored regions, % | | 88.21 | 90.23 | 88.31 |
| Allowed regions, % | | 9.55 | 7.97 | 9.53 |
| Outliers, % | | 2.24 | 1.80 | 2.16 |
| *Deviations from ideal values (RMSD)* | | | | |
| Bond, Å | | 0.010 | 0.010 | 0.009 |
| Angle, degrees | | 1.581 | 1.639 | 1.519 |
| Chirality | | 0.065 | 0.067 | 0.065 |
| Planarity | | 0.008 | 0.008 | 0.008 |
| Dihedral, degrees | | 17.572 | 17.157 | 17.391 |
| Average B-factor (overall), Å$^2$ | | 54.5 | 57.9 | 56.2 |

**Table S1 (continued). X-ray data collection and refinement statistics.**

| ***Crystals*** | | **70S-PY complex with ACCA-ITM, ACCA, and CHL** | **70S-PY complex with ACCA-IAM, ACCA, and CHL** |
|---|---|---|---|
| ***Diffraction data*** | | | |
| Space Group | | $P2_12_12_1$ | $P2_12_12_1$ |
| Unit Cell Dimensions, Å (a x b x c) | | 209.71 x 449.95 x 621.20 | 209.81 x 449.56 x 621.47 |
| Wavelength, Å | | 0.9795 | 0.9795 |
| Resolution range (outer shell), Å | | 310-2.50 (2.56-2.50) | 311-2.40 (2.46-2.40) |
| $I/\sigma I$ (outer shell) | | 8.41(0.85) | 7.95 (0.85) |
| Resolution at which $I/\sigma I$=1, Å | | 2.50 | 2.40 |
| Resolution at which $I/\sigma I$=2, Å | | 2.75 | 2.65 |
| CC(1/2) at which $I/\sigma I$=1, % | | 16.0 | 10.0 |
| CC(1/2) at which $I/\sigma I$=2, % | | 50.0 | 40.0 |
| Completeness (outer shell), % | | 98.5 (93.1) | 99.8 (99.4) |
| $R_{merge}$ (outer shell)% | | 16.0 (147.2) | 24.6 (234.2) |
| No. of crystals used | | 1 | 2 |
| No. of Reflections Used: | Observed | 9,083,103 | 20,837,902 |
| | Unique | 1,966,210 | 2,250,029 |
| Redundancy (outer shell) | | 4.62 (3.87) | 9.26 (8.80) |
| ***Refinement*** | | | |
| Resolution range of the diffraction data included in the refinement, Å | | 225-2.50 | 211-2.40 |
| $R_{work}/R_{free}$, % | | 22.5/28.3 | 22.1/27.2 |
| *No. of Non-Hydrogen Atoms* | | | |
| RNA | | 192,599 | 192,599 |
| Protein | | 93,149 | 93,145 |
| Ions (Mg, K, Zn, Fe) | | 2,548 | 2,548 |
| Waters | | 9,495 | 9,498 |
| *Ramachandran Plot* | | | |
| Favored regions, % | | 87.08 | 88.76 |
| Allowed regions, % | | 10.32 | 9.01 |
| Outliers, % | | 2.60 | 2.23 |
| *Deviations from ideal values (RMSD)* | | | |
| Bond, Å | | 0.009 | 0.009 |
| Angle, degrees | | 1.467 | 1.459 |
| Chirality | | 0.061 | 0.061 |
| Planarity | | 0.007 | 0.007 |
| Dihedral, degrees | | 17.703 | 17.393 |
| Average B-factor (overall), $Å^2$ | | 54.4 | 55.1 |

#### III. SUPPLEMENTARY FIGURES

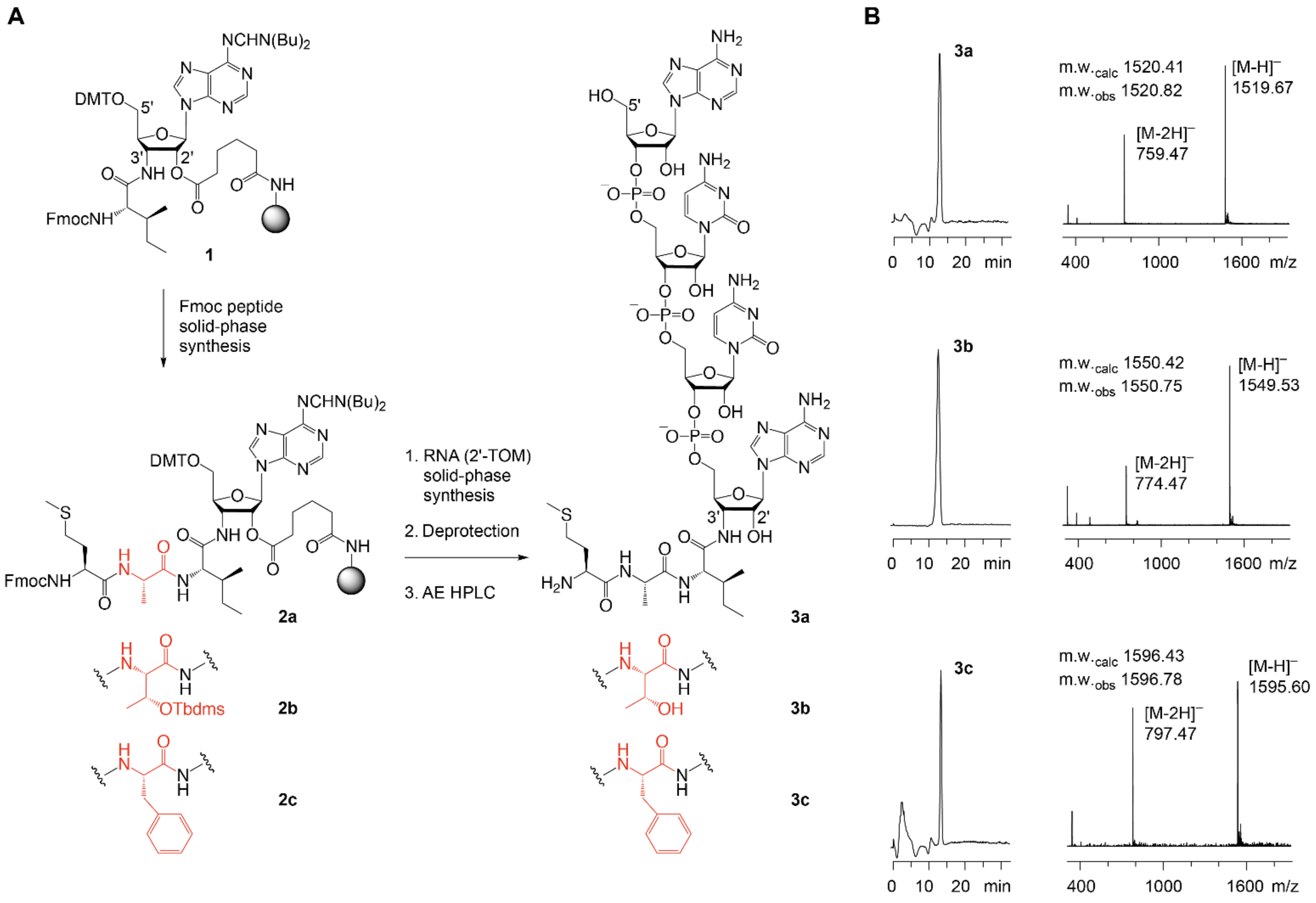


**Figure S1. Chemical synthesis of hydrolysis-resistant tripeptidyl-ACCA conjugates.** (**A**) Overview of the synthetic pathway. Chemical structure of functionalized solid support **1**[2] (grey sphere represents amino-functionalized polystyrene support (GE Healthcare, Custom Primer SupportTM 200 Amino) used for peptide assembly (Fmoc chemistry) and RNA assembly (2'-O-TOM chemistry), followed by deprotection and purification using anion-exchange chromatography; DMT = 4,4'-dimethoxytrityl, Fmoc = N-(9-fluorenyl)methoxy-carbonyl, TOM [(triisopropylsilyl)oxy]methyl, AE HPLC = anion-exchange high-pressure liquid chromatography. (**B**) Anion-exchange HPLC profiles of purified ACCA-Ile-Ala-Met, ACCA-Ile-Thr-Met, and ACCA-Ile-Phe-Met conjugates (left) and LC-ESI mass spectra (right). Anion-exchange chromatography conditions: Dionex DNAPac PA-100 (4x250 mm) column; temperature: Flow rate: 1 mL/min; eluent A: 25 mM Tris-HCl (pH 8.0) and 20 mM $NaClO_4$ in 20% aqueous acetonitrile, eluent B: 25 mM Tris-HCl (pH 8.0) and 0.60 M $NaClO_4$ in 20% aqueous acetonitrile; gradient: 0-35% B in A within 28 min; UV detection at $\lambda$ = 260 nm.

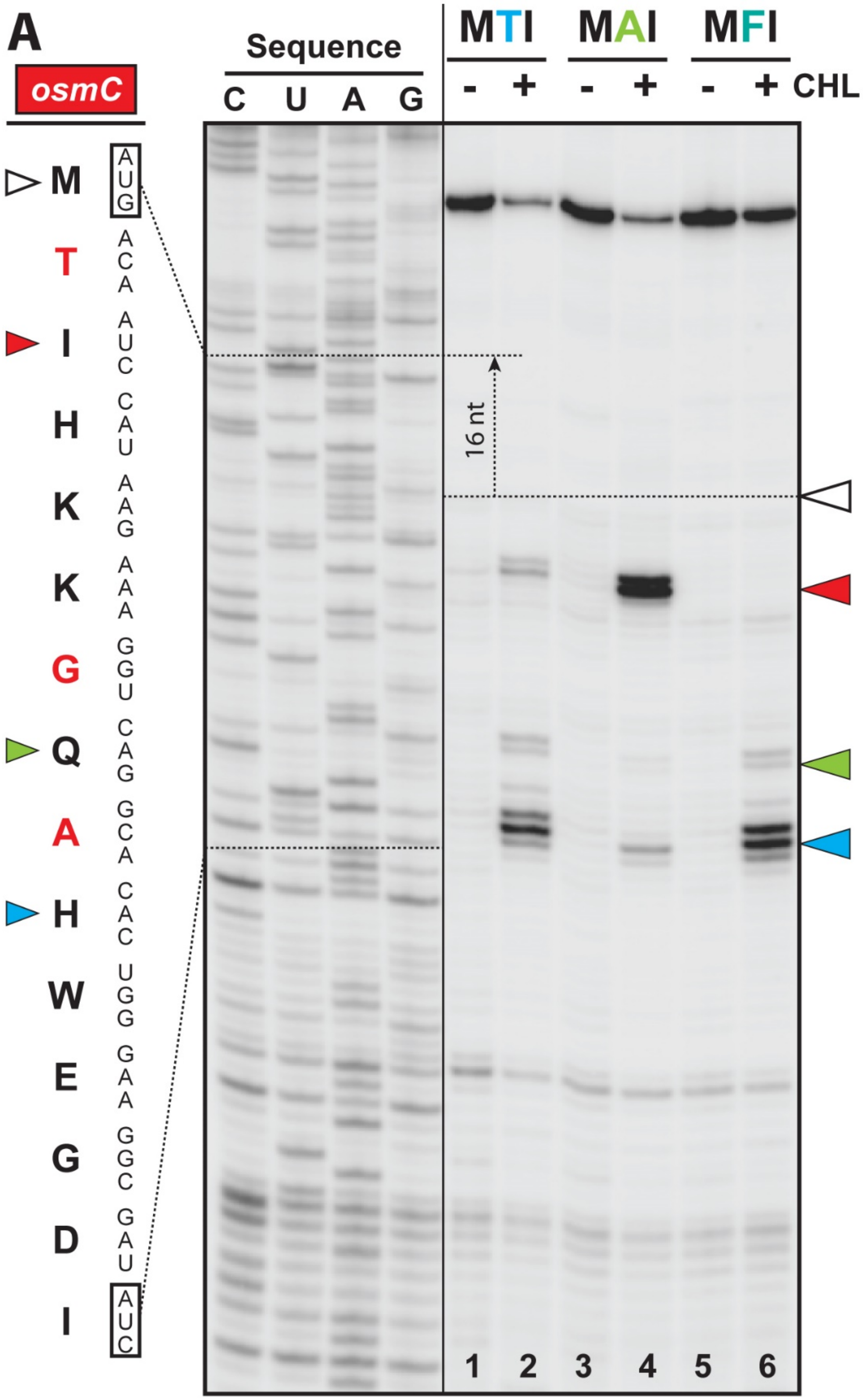


**Figure S2. CHL arrests translation at the MTI/MAI tripeptide sequences.** Ribosome stalling in the presence and absence of CHL revealed by reverse-transcription primer-extension inhibition (toe-printing) assay in a cell-free translation system on wild-type *osmC* mRNA encoding MTI tripeptide at the Nterminus (lanes 1-2) or its mutant versions encoding either alanine (lanes 3-4) or phenylalanine (lanes 5-6) in the 2$^{nd}$ position. Nucleotide sequences of wild-type *osmC* mRNA and the corresponding amino acid sequence are shown on the left. White arrowhead marks translation start site. Red and blue arrowheads point to the drug-induced arrest sites within the coding sequences of each of the three used mRNAs. Note that due to the large size of the ribosome, the reverse transcriptase used in the toe-printing assay stops 16 nucleotides downstream of the codon located in the P-site. Because *osmC* mRNA template harbors other downstream alanine codons, CHL also caused ribosome stalling at these downstream sites subsequent to the appearance of Ala, Ser, or Thr residues in the penultimate position of the peptide chain (lanes 2, 4, and 6, blue arrowhead). The results of the toe-printing assay confirmed that the presence of the MTI or MAI tripeptide sequences in the P site of the ribosome results in CHL-dependent stalling, whereas MFI tripeptide sequence is not conducive to ribosome stalling and can serve as a negative control.

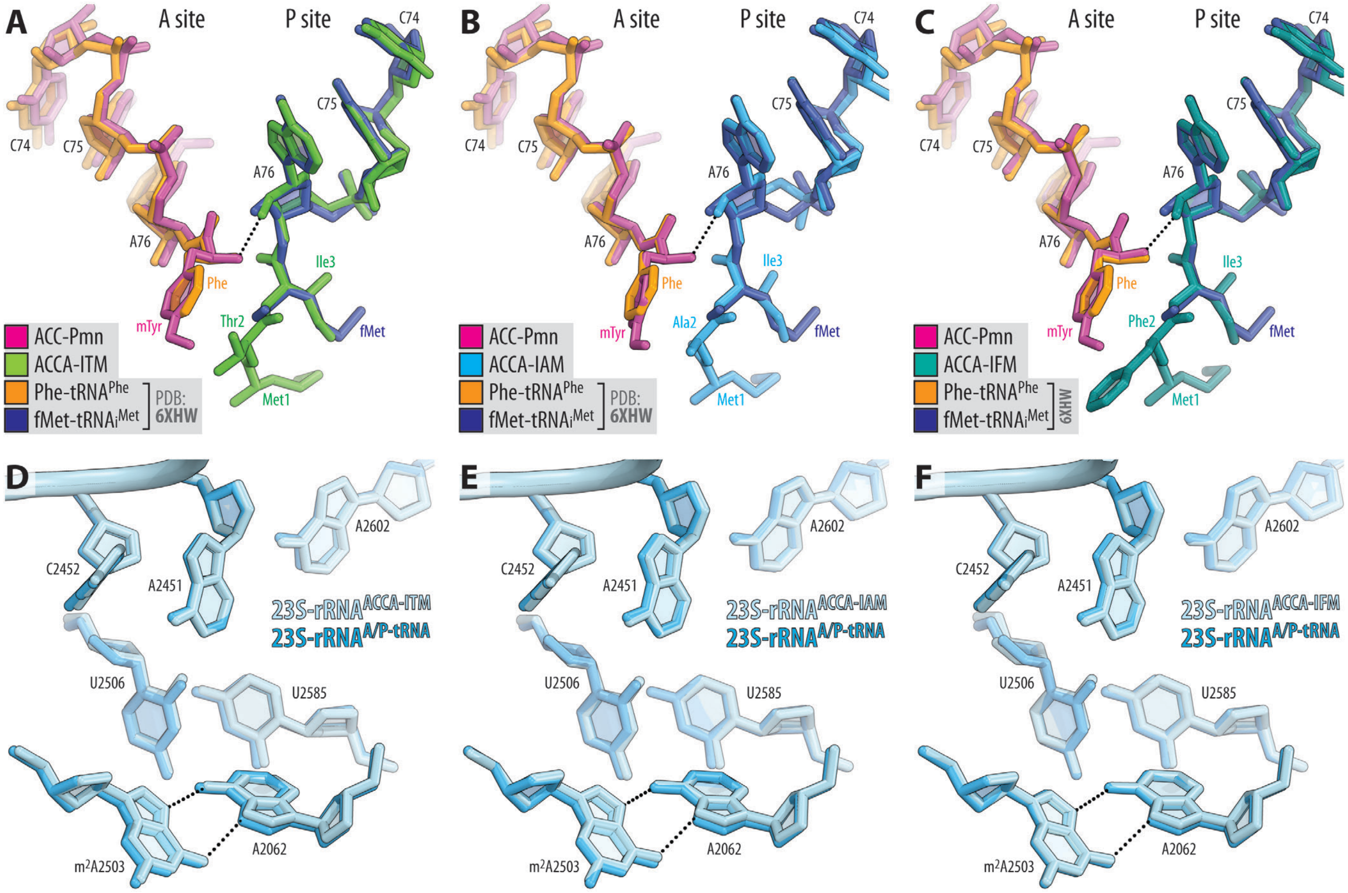


**Figure S3. Superpositioning of the structures of short tRNA analogs with the aminoacylated full-length tRNAs.** (**A, B, C**) Comparisons of the 70S ribosome structures carrying ACCA-PMN (magenta) in the A site and either ACCA-ITM (A, green), or ACCA-IAM (B, blue), or ACCA-IFM (teal) short peptidyl tRNA analogs in the P site with the previous structure of ribosome-bound full-length aminoacyl-tRNAs (PDB entry 6ZHW[15]). All structures were aligned based on domain V of the 23S rRNA. (**D, E, F**) Comparisons of the positions of key 23S rRNA nucleotides around the PTC in the same structures. Note that there are no significant differences in the positions of A- or P-site substrates or the PTC nucleotides.

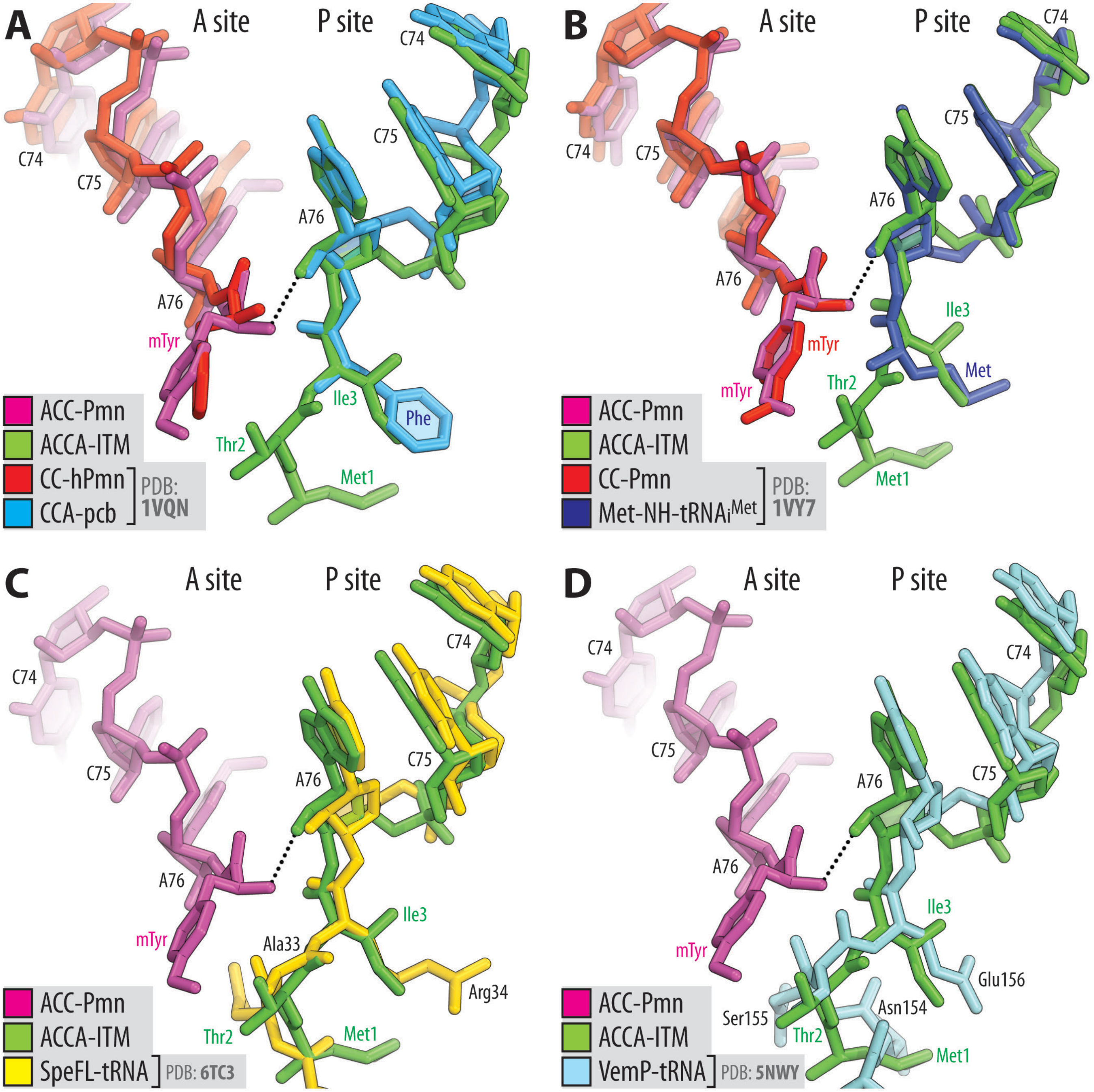


**Figure S4. Comparison of the new structure of short MTI-tripeptidyl-tRNA analog with other structures of ribosome-bound short tRNA analogs and stalling peptides.**

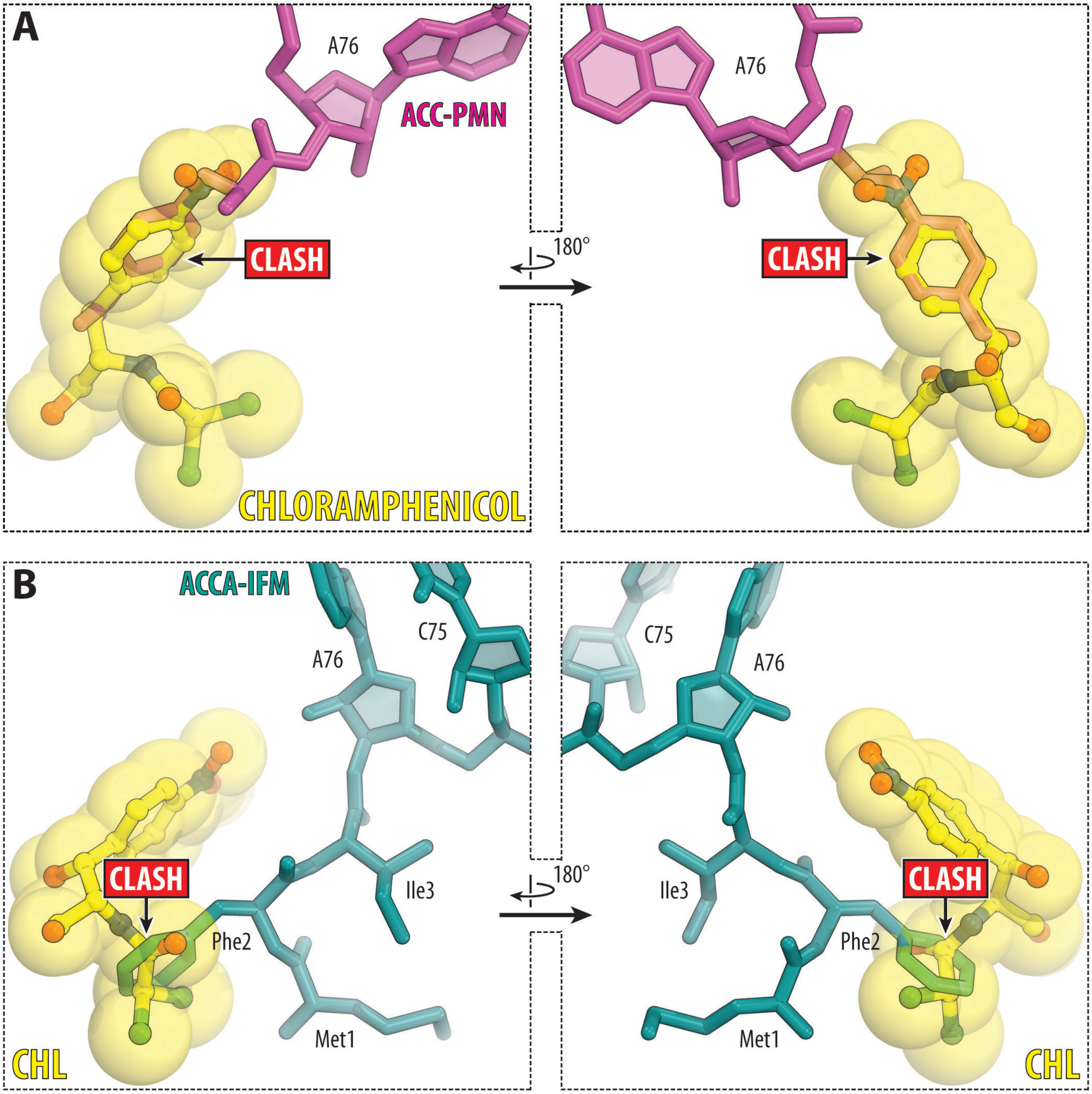


**Figure S5. Superposition of the ribosome-bound CHL with the structures of ribosome-bound aa-tRNA amd peptidyl-tRNA analogs.** (**A**) Superposition of CHL with the A-site-bound aa-tRNA analog ACC-PMN. (**B**) Superposition of CHL with the P-site-bound peptidyl-tRNA analog ACCA-IFM. The structure of CHL is from PDB entry 6ND5[14]. The structures were aligned based on domain V of the 23S rRNA. Note that the side chains of the incoming amino acid in the A site (A) or the penultimate amino acid of the peptide in the P site (B) clash with CHL in its canonical binding site.

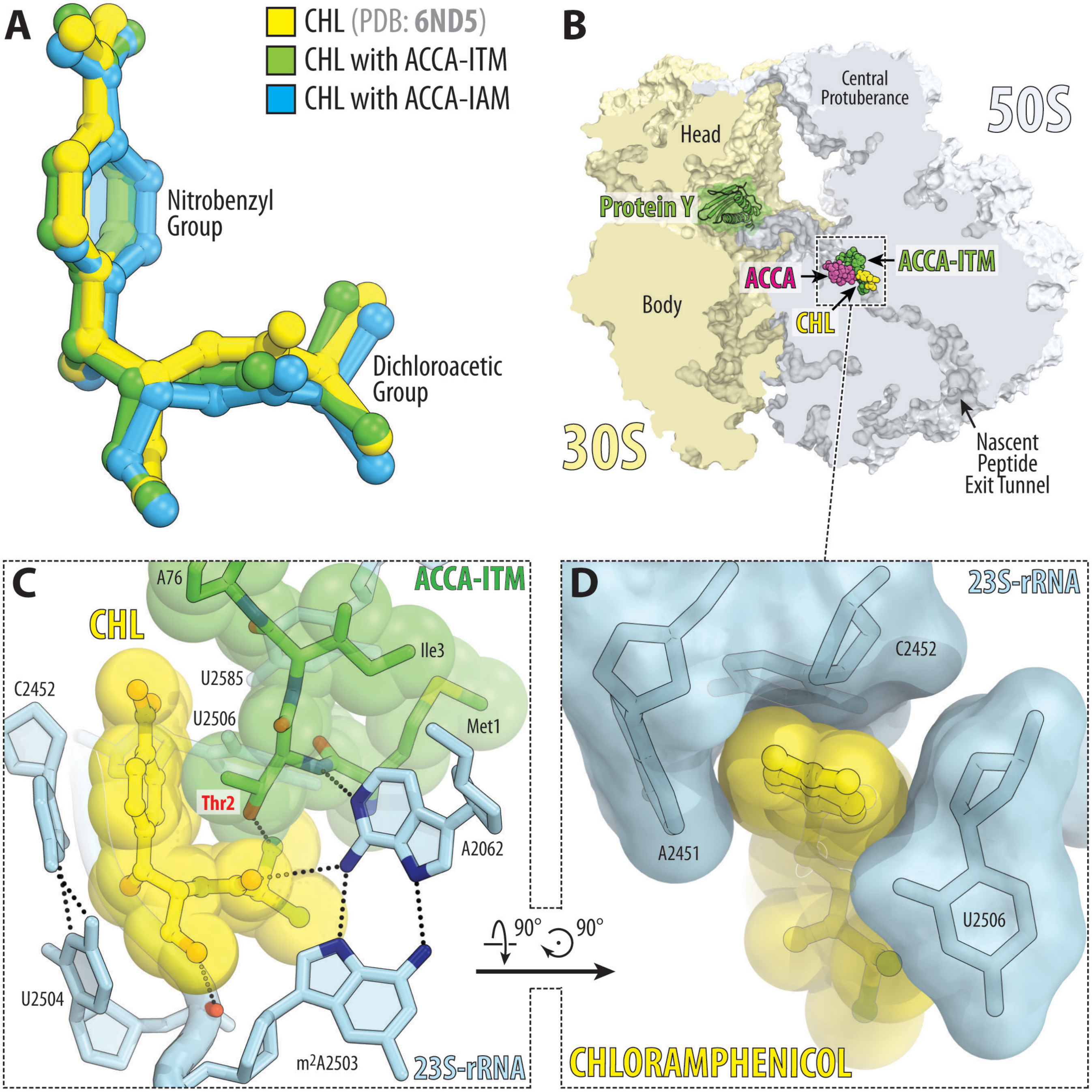


**Figure S6. Structure of CHL in complex with the 70S ribosome and MTI-tripeptidyl-tRNA analog.** (**A**) Superposition of the previous structure of CHL bound to the 70S ribosome in the presence of deacylated A- and P-site tRNAs (yellow, PDB entry 6ND5[14]) with the new structures of CHL in the presence of MTI (green) or MAI (blue) tripeptides. All structures were aligned based on domain V of the 23S rRNA. (**B**) Overview of the CHL binding site (yellow) in the *T. thermophilus* 70S ribosome in complex with the short tripeptidyl-tRNA analogs viewed as a cross-cut section through the nascent peptide exit tunnel. The 30S subunit is shown in light yellow; the 50S subunit is in light blue; ribosome-bound protein Y is colored in green. (**C, D**) Close-up views of CHL bound in the PTC, highlighting H-bond interactions (dashed lines) and the intercalation of the nitrobenzyl group into the A-site cleft formed by nucleotides A2451 and C2452 of the 23S rRNA. Note that the side chain of the Thr2 residue of the MTI tripeptide directly interacts with the ribosome-bound CHL.

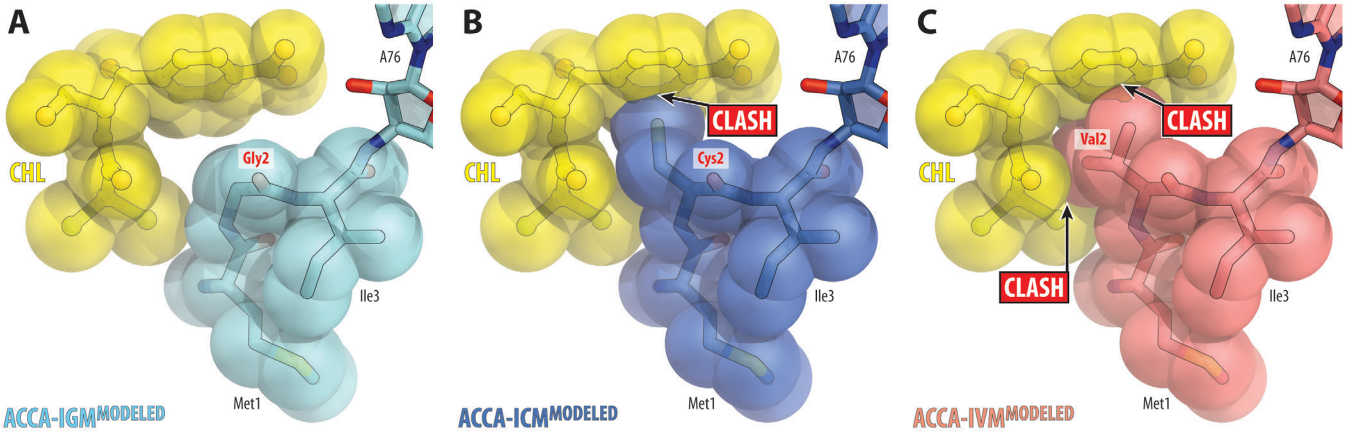


**Figure S7. *In silico* modeling of MGI, MCI, and MVI tripeptides in the presence of CHL.** Using the structure of MAI-tripeptidyl-tRNA analog bound to the ribosome in the presence of CHL as a reference, we mutated the second alanine residue either to glycine (**A**), cysteine (**B**), or valine (**C**) and assessed the modeled structures for sterical clashes.

#### IV. SUPPLEMENTARY REFERENCES


1. Steger, J. et al. Efficient access to nonhydrolyzable initiator tRNA based on the synthesis of 3'-azido-3'-deoxyadenosine RNA. *Angew. Chem. Int. Ed. Engl.* **49**, 7470-7472 (2010).

2. Steger, J. & Micura, R. Functionalized polystyrene supports for solid-phase synthesis of glycyl-, alanyl-, and isoleucyl-RNA conjugates as hydrolysis-resistant mimics of peptidyl-tRNAs. *Bioorg. Med. Chem.* **19**, 5167-5174 (2011).

3. Geiermann, A.S., Polacek, N. & Micura, R. Native chemical ligation of hydrolysis-resistant 3'-peptidyl-tRNA mimics. *J. Am. Chem. Soc.* **133**, 19068-19071 (2011).

4. Moroder, H. et al. Non-hydrolyzable RNA-peptide conjugates: a powerful advance in the synthesis of mimics for 3'-peptidyl tRNA termini. *Angew. Chem. Int. Ed. Engl.* **48**, 4056-4060 (2009).

5. Marks, J. et al. Context-specific inhibition of translation by ribosomal antibiotics targeting the peptidyl transferase center. *Proc. Natl. Acad. Sci. USA* **113**, 12150-12155 (2016).

6. Orelle, C. et al. Tools for characterizing bacterial protein synthesis inhibitors. *Antimicrob. Agents Chemother.* **57**, 5994-6004 (2013).

7. Polikanov, Y.S., Steitz, T.A. & Innis, C.A. A proton wire to couple aminoacyl-tRNA accommodation and peptide-bond formation on the ribosome. *Nat. Struct. Mol. Biol.* **21**, 787-793 (2014).

8. Melnikov, S.V. et al. Mechanistic insights into the slow peptide bond formation with D-amino acids in the ribosomal active site. *Nucleic Acids Res.* **47**, 2089-2100 (2019).

9. Polikanov, Y.S., Blaha, G.M. & Steitz, T.A. How hibernation factors RMF, HPF, and YfiA turn off protein synthesis. *Science* **336**, 915-918 (2012).

10. Polikanov, Y.S., Melnikov, S.V., Soll, D. & Steitz, T.A. Structural insights into the role of rRNA modifications in protein synthesis and ribosome assembly. *Nat. Struct. Mol. Biol.* **22**, 342-344 (2015).

11. Tereshchenkov, A.G. et al. Binding and action of amino acid analogs of chloramphenicol upon the bacterial ribosome. *J. Mol. Biol.* **430**, 842-852 (2018).

12. Chen, C.W. et al. Binding and action of triphenylphosphonium analog of chloramphenicol upon the bacterial ribosome. *Antibiotics (Basel)* **10**(2021).

13. Almutairi, M.M. et al. Co-produced natural ketolides methymycin and pikromycin inhibit bacterial growth by preventing synthesis of a limited number of proteins *Nucleic Acids Res.* (2017).

14. Svetlov, M.S. et al. High-resolution crystal structures of ribosome-bound chloramphenicol and erythromycin provide the ultimate basis for their competition. *RNA* **25**, 600-606 (2019).

15. Svetlov, M.S. et al. Structure of Erm-modified 70S ribosome reveals the mechanism of macrolide resistance. *Nat. Chem. Biol.* **17**, 412-420 (2021).

16. Schuttelkopf, A.W. & van Aalten, D.M. PRODRG: a tool for high-throughput crystallography of protein-ligand complexes. *Acta Crystallogr. D Biol. Crystallogr.* **60**, 1355-1363 (2004).

17. Herrero Del Valle, A. et al. Ornithine capture by a translating ribosome controls bacterial polyamine synthesis. *Nat. Microbiol.* **5**, 554-561 (2020).

18. Su, T. et al. The force-sensing peptide VemP employs extreme compaction and secondary structure formation to induce ribosomal stalling. *Elife* **6**(2017).

19. Goto, Y. & Suga, H. Translation initiation with initiator tRNA charged with exotic peptides. *J. Am. Chem. Soc.* **131**, 5040-5041 (2009).
